# Tensor-Based Integration of Time-Series Measurements Reveals Relationships Between Underlying Disease, Early Injury, and CD4+ Polarization in Liver Transplantation

**DOI:** 10.1101/2025.05.29.655766

**Authors:** Jackson L. Chin, Zhixin Cyrillus Tan, Rebecca A. Sosa, Ying Zheng, David W. Gjertson, Alexander Hoffmann, Fady M. Kaldas, Yuan Zhai, Jerzy W. Kupiec-Weglinski, Elaine F. Reed, Aaron S. Meyer

## Abstract

For patients with end-stage liver failure, liver transplantation (LT) remains the standard-of-care, though five-year allograft survival rates remain below 80%. Liver ischemia reperfusion injuries (LIRI) arise during transplant and contribute to allograft dysfunction. While many drivers of LIRI have been well-characterized, the relationships between LIRI and later immunological signatures remain poorly understood, possibly because of limited integrated studies examining immunological signatures across the LT process. Furthermore, the role of underlying disease in LT remains poorly understood as it is unknown if later immunological signatures are universal or etiology dependent. To better examine etiology- and LIRI-driven immunological mechanisms across the LT lifetime, we collected cytokine and liver function test (LFT) measurements from LT donors and recipients at multiple pre- and post-operative timepoints. By developing a tensor-based factorization method to integrate these longitudinal measurements, we reduced the datasets to four immunological signatures that manifested across cytokines, LFTs, and time. Correlative analyses revealed associations between these integrated immunological signatures with etiology and LIRI. Later integration of allograft survival into this factorization method reduced these four signatures to two that were predictive of five-year allograft survival. Examination of these signatures prognostic of allograft survival found that CD4+ hyperpolarization towards Th1 or Th2 subtypes impaired growth factor expression and led to increased allograft loss risk. Clinical correlates revealed that donor and recipient age alongside etiology and donor liver health modulate this balance of CD4+ polarization in LT. Collectively, these results link immunological mechanisms to etiology and LIRI offering immediately actionable clinical insights alongside etiology- and donor-specific therapeutic targets for improving LT outcomes.

## Introduction

For patients with end-stage liver disease, liver transplantation (LT)—wherein the patient’s liver is replaced with either a portion of a healthy liver from a living donor or a complete liver from a deceased donor— remains the standard-of-care^1,2^. A myriad of underlying etiologies can cause end-stage liver disease, including dietary-based diseases like alcoholic liver disease (ALD), chronic diseases like nonalcoholic steatohepatitis (NASH), infectious diseases such as hepatitis C virus (HCV), and cancers like hepatocellular carcinoma (HCC)^3^. These diseases induce irreversible scarring, or cirrhosis, in the liver that impair function and can eventually progress to end-stage disease that requires a transplant to remedy.

Despite their clinical importance and high demand, transplanted livers and their recipients exhibit poor long-term survival rates; one-year survival rates for patients and allografts are at 94.9% and 93.6%, respectively, decreasing to 81.2% and 78.4% five years post-operation^4^. A common contributor to patient and allograft loss in LT is early allograft dysfunction (EAD) characterized by impaired post-transplant liver function and elevated liver damage in the first post-operative week^5^. EAD can impair patient quality-of-life as LT recipients with EAD typically have longer ICU stays than their non-EAD counterparts. Furthermore, EAD negatively impacts both short- and long-term outcomes, reducing patient and allograft survival rates at 1-, 3-, and 5-years^6,7^.

Liver ischemia reperfusion injuries (LIRI) accrued across the transplant process are thought to contribute to EAD^8,9^ and are often divided between two phases: cold and warm ischemic reperfusion injuries. Cold ischemia injuries are accrued while the donor liver is cooled and transported following organ harvest and consist predominantly of ischemia-related damage, such as cell death, reactive oxygen species (ROS) formation, and mitochondrial dysfunction^10–12^. Hepatocellular damage from these injuries releases damage-associated molecular patterns (DAMPs) that exacerbate future injuries, such as those in warm ischemic phases. Warm ischemia injuries, conversely, consist of injuries incurred after cold transit and frequently involve both donor and recipient immune responses. Early warm ischemia reperfusion injuries follow anastomosis and involve innate inflammatory responses driven by liver-resident Kupffer cells (KCs)^13–15^ alongside natural killer (NK) cells from both the donor and recipient^16,17^. Such early warm ischemia responses later give way to granulocyte- and macrophage-mediated inflammation as well as adaptive immune responses that can exacerbate hepatocellular damage and modulate immune responses to LT^14,18–22^.

While prior efforts have found many drivers of LIRI, the relationships between temporally disparate LIRI sources and their connection to liver failure etiology and LT outcome remain poorly understood. We hypothesize that integration of longitudinal donor and recipient measurements collected across the LT process will provide further clarity in the relationships between injury sources in LT. We posit that integration of longitudinal immunological features will capture undiscovered relationships between temporally disparate LIRI sources. Furthermore, as both donor and recipient factors drive LIRI, integration of recipient and donor measurements would more comprehensively characterize immunological signatures across time and tissues for each patient, thus improving our ability to understand the relationships between LIRI sources, etiology, and eventual LT outcome.

To explore this hypothesis, we collected whole-blood cytokine measurements from both recipient and donor tissues at multiple pre- and post-operative time points alongside recipient liver function test (LFT) measurements. We then apply tensor-based factorization to integrate these longitudinal molecular and clinical measurements to capture temporally evolving immune signatures. Through interpretation of this factorization model, we find relationships between LIRI drivers and etiology seen across the LT process. Later, by incorporating allograft loss outcomes into our factorization, we refine these four processes to two critical immunological patterns predictive of LT outcome. Comparisons to single dataset and clinical data models reveal that integrative and outcome-informed factorization improves prognosis of LT outcome while identifying critical immunological signatures determining LT fate. These findings help to characterize the relationships between temporally disparate LIRI mechanisms, etiology, and transplant outcome.

## Results

### Longitudinal Clinical & Molecular Measurements Capture Transient, Etiology- and LIRI-Driven Immunological Perturbations

To comprehensively characterize immunological signatures across the LT process, we collected three longitudinal datasets from a series of LTs (n=144) performed at a single center (Figure 1A). Cytokine measurements were collected from the peripheral blood of LT recipients pre-operatively (PO) and at one-day (D1), one-week (W1), and one-month (M1) post-operation (Figure 1A–B). Three liver function tests (LFTs)—bilirubin (TBIL), alanine transaminase (ALT), and aspartate transaminase (AST)—were collected pre-operatively and daily for the first week post-operation to quantify specific types of liver damage (Figure 1B). Finally, to characterize donor immune signatures, cytokine measurements were collected from recipient intraoperative portal vein (PV) blood taken prior to reperfusion (PR) and during reperfusion as recipient PV blood was flushed through the donor vena cava (liver flush, LF) (Figure 1B). A small subset of LTs were missing specific measurements at certain timepoints (Figure S1).

**Figure 1:**
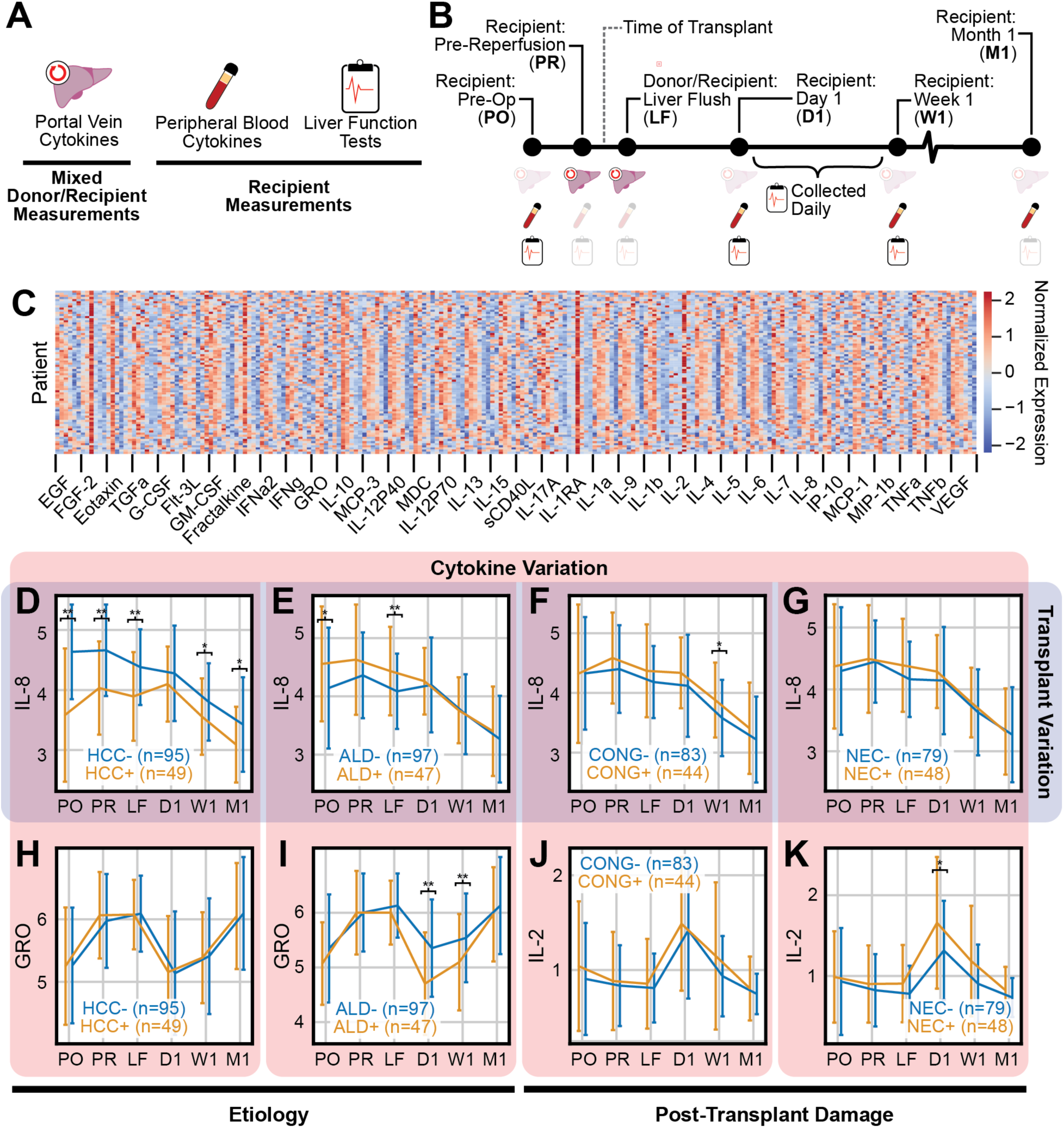
Longitudinal Measurements Capture Transient Variation in Immunological Signatures Driven by Etiology and LIRI. A) Overview of measurements; cytokine measurements and liver function tests (LFT) were collected from peripheral blood of liver transplant recipients alongside cytokine measurements from the portal vein of donor livers. B) Measurement and sample collection timeline; (top) samples were collected at multiple timepoints across the liver transplant process; (bottom) each measurement type was collected at different timepoints, though each measurement was collected at least once pre-operatively and once post-operatively. C) Heatmap of cytokine measurements, where each row corresponds to a patient and each column represents a cytokine measurement at a specific timepoint. Each bin corresponds to a different cytokine; columns within each bin are ordered chronologically from left to right. Measurements were normalized for each cytokine across timepoints to have zero mean and unit variance. D–K) Line plots depicting arithmetic mean of IL-8 (D–G), GRO (H–I), and IL-2 (J–K) expression at each measurement timepoint; error bars correspond to standard deviation in expression. Patients are subset by presence of hepatocellular cancer (D/H), alcoholic liver disease (E/I), post-operative sinusoidal congestion (F/J), and post-operative necrosis (G/K). Cytokine expression is compared at each timepoint via independent *t*-test. * denotes *p*<0.05, ** denotes *p*<0.01 level.

Longitudinal measurements may offer new insights into the dynamic immunological processes underpinning LT. Preliminary examination of the cytokine measurements revealed that expression was unique across time, transplant, and cytokine (Figure 1C–K). For example, widespread temporal variation was observed in the heatmap of cytokine measurements (Figure 1C); vertical bands of color show systemic increases (red) or decreases (blue) in cytokine expression at a timepoint across patients.

Variation in expression also manifested uniquely across cytokines based on both recipient and donor factors. For instance, IL-8 temporal dynamics differed by etiology (Figure 1D–E) and with post-transplant markers of LIRI (Figure 1F–G). LT recipients with hepatocellular cancer (HCC) exhibited universally lower IL-8 (Figure 1D) while those with alcoholic liver disease (ALD) showed higher IL-8 pre-operatively and shortly after transplant (Figure 1E). Likewise, IL-8 expression was higher at D1 in LTs with post-transplant sinusoidal congestion (Figure 1F) and was independent of post-transplant necrosis (Figure 1G). These temporal variations were not uniform across different cytokines. For instance, GRO expression was independent of HCC status (Figure 1H) and was reduced post-operation for recipients with ALD (Figure 1I); IL-2 expression was independent of post-transplant sinusoidal congestion (Figure 1J) but was elevated at D1 in LTs with post-transplant necrosis (Figure 1K). The LFT measurements demonstrated similar transplant-specific temporal variation (Figure S2). LFTs elevated above normal ranges are typical markers for EAD^5^; within the first post-operative week, however, the timing of this elevation varied across transplants. Most patients had elevated LFTs at the time-of-transplant, but many experienced a decrease in their scores that rebounded later in the first week after surgery (Figure S2B–D). Moreover, these temporal patterns were not uniform across different LFT types.

### Integration of Longitudinal Measurements Yields Four Distinct Integrative Immune Signatures in LT Transplant

While longitudinal measurements offer insights into temporally evolving immunological signatures, a challenge with such datasets is in separating patient-specific from longitudinal patterns. Each dataset consists of three biological dimensions that each drive variation in measurements: patients, time, and cytokines (peripheral and PV cytokine datasets) or LFT scores (LFT dataset).

To most effectively integrate these longitudinal measurements and separate variation seen across dimensions, we restructured each dataset into a tensor. Tensors are organized arrays with three or more dimensions (modes)^23^. In this study, each of the three longitudinal datasets (peripheral/PV cytokines, LFTs) were organized into three-mode tensors by “stacking” measurements collected at each timepoint to form a cube. For example, the peripheral cytokine measurements were represented as a series of expression matrices corresponding to patient cytokine measurements at each sampling timepoint (Figure 2A). These matrices were then stacked as they share a common set of patients and cytokines, producing a three-mode tensor with patient, time, and cytokine dimensions. Tensor-based factorization techniques then reduce tensor-structured datasets to a set of “components” that capture patterns observed across dimensions (Figure 2B). These components were interpreted through factor matrices; tensor factorization methods produce one factor matrix for each dimension, and each factor matrix defines relationships between each component pattern and its corresponding biological dimension (Figure 2C). When appropriate, tensor-based approaches often improve the effectiveness of data reduction^23^, and the resulting factor matrices help to better separate and visualize variation observed within each dimension^24–27^.

**Figure 2:**
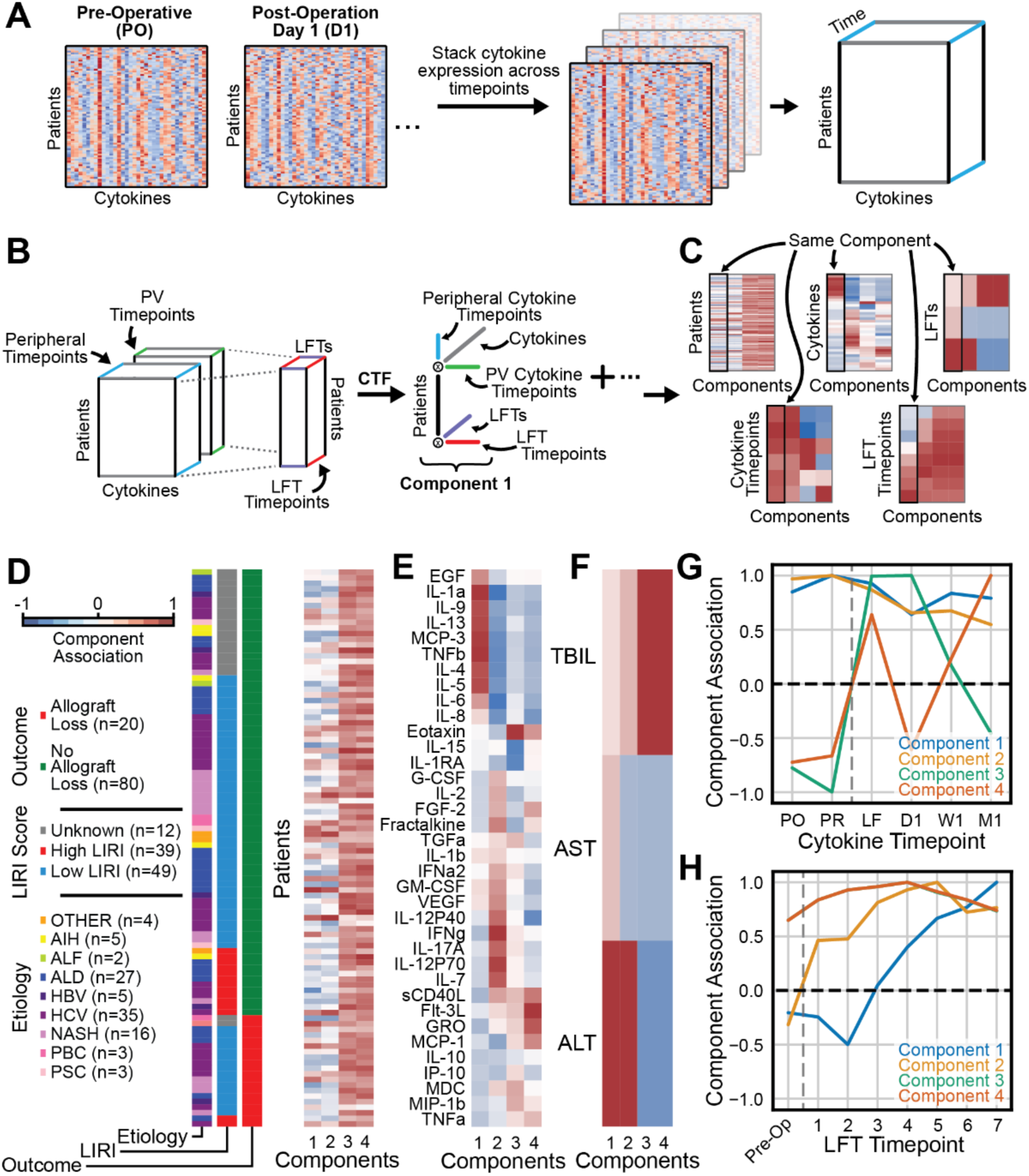
Integration of Cytokine and LFT Measurements Captures Longitudinal Relationships Among Immunological Signatures and Damage Patterns. A) Overview of tensor setup. Longitudinal measurements, like the cytokine measurements, can be represented as a series of expression matrices. As patients and cytokines are collected consistently across timepoints, these expression matrices can be stacked into a three-dimensional tensor, or cube, with dimensions of patients, time, and cytokines. B) Each of the three datasets (peripheral/PV cytokines, LFTs) were restructured into three-dimensional tensors; as each dataset consists of the same patients, the patient dimension can be coupled across datasets. Similarly, as the same cytokines are assayed in the peripheral and PV cytokine datasets, the cytokine dimension between these tensors was also coupled. Coupled tensor factorization (CTF) can then reduce these coupled tensors to a set of components observed across longitudinal datasets. C) Factor matrices produced via CTF can then be used to biologically interpret each component. CTF produces five factor matrices, one for each dimension in the coupled datasets, that define relationships between each respective biological dimension and the CTF components. D–F) Patient (D), cytokine (E), and LFT (F) factor matrices for a four component CTF decomposition. Bars in (D) denote transplant outcome, LIRI score, and primary end stage liver failure etiology. Abbreviations correspond to the following etiologies: autoimmune hepatitis (AIH), acute liver failure (ALF), alcoholic liver disease (ALD), hepatitis B virus (HBV), hepatitis C virus (HCV), non-alcoholic steatohepatitis (NASH), primary biliary cirrhosis (PBC), and primary sclerosing cholangitis (PSC). G–H) Line plots depicting component associations to cytokine measurement timepoints (G) and LFT measurement timepoints (H). Timepoints are in chronological order from left to right. Vertical dashed line corresponds to time-of-transplant.

To capture variation observed across longitudinal datasets, we applied a variant of tensor factorization, coupled tensor factorization (CTF), to integrate the peripheral cytokine, PV cytokine, and LFT tensors (Figure 2B). Like traditional tensor factorization methods, CTF reduces each dataset tensor to a set of components; however, CTF recognizes that each tensor comprises the same group of patients and couples the patient factor across tensors by sharing a common patient factor in each tensor’s factorization. Similarly, as the same cytokines are assayed in both the peripheral and PV cytokine datasets, the cytokine dimension is also coupled in CTF. As these dimensions are coupled, CTF should therefore capture patterns observed across datasets, and we hypothesized that this approach may most effectively integrate longitudinal measurements.

Factorization via CTF requires the user to specify the number of components used in the resulting factorization. To choose this number of components, we applied CTF to resampled sets of patients and evaluated the consistency of the resulting factor matrices via Factor Match Score (Figure S3). The CTF components were most consistent when two or four components were used (Figure S3A–B), and we ultimately chose to use the four-component CTF as it captured more information in the resulting factorization (Figure S3C). CTF captured a similar amount of variance compared to principal component analysis (PCA) applied to a matrix form of the longitudinal datasets, indicating that CTF’s information capture was similar to other common dimensionality reduction techniques (Figure S3C).

### CTF Links Recurrent Th2 Responses with Donor Liver Health and Identifies Protective Eotaxin Responses Reduced in Patients with Alcoholic Liver Disease

To interpret the biological significance of the CTF model, we plotted the five resulting factor matrices: 1) patients, 2) cytokines, 3) LFTs, 4) cytokine measurement timepoints, and 5) LFT measurement timepoints (Figure 2D–H). As measurement timepoints are ordered, the cytokine and LFT measurement timepoint factor matrices were plotted as line plots (Figure 2G–H); timepoints are in chronological order, and the PV and peripheral cytokine measurements were combined into one axis. These factor matrices can be interpreted in two primary ways to identify either (1) intra-dimensional variation or (2) integrated, multidimensional biological patterns.

To evaluate intra-dimensional variation, one can examine differences in component associations within each dimension. For instance, CTF 1 and 2 were similar in their associations to LFT scores as both corresponded to elevated ALT (Figure 2F); CTF 1 and 2 were unique, however, in that CTF 1 corresponded to ALT elevation later post-operation (post-operation day 4) than CTF 2 (post-operation day 1) (Figure 2H). CTF 1 and 2 also demonstrated similar associations to cytokine timepoints as both corresponded to cytokines that were most elevated pre-operatively at the PO and PR measurements before attenuating at the D1 timepoint (Figure 2G); CTF 1 and 2, however, corresponded to opposing groups of cytokines as many of the cytokines elevated in CTF 1—like IL-1a, IL-4, IL-5, IL-9, and IL-13— were reduced by the signatures represented in CTF 2 (Figure 2E).

Alternatively, one can also trace a component across matrices to examine immunological patterns observed across time and datasets. For example, CTF 1 associated with IL-1a, IL-9, IL-13, MCP-3, TNFβ, IL-4, and IL-5 (Figure 2E) that were elevated at most measurement timepoints with a dip in magnitude at the D1 timepoint (Figure 2G). In conjunction, this cytokine signature also associated with ALT (Figure 2F) that was initially reduced before elevating later post-operation (Figure 2H). These observations link this recurring cytokine signature with a delayed elevation in ALT-associated liver damage.

To corroborate the recurring cytokine signature indicated by this component, we compared pre-operative expression of cytokines associated with CTF 1—including Th2 cytokines like IL-4, IL-5, IL-9, and IL-13— with their expression at the D1, W1, and M1 timepoints. Indeed, pre-operative expression of these cytokines was significantly correlated with their post-operative expression (Figure 3A–C). Furthermore, pre-operative expression of these CTF 1 cytokines was more strongly correlated with later post-operative timepoints (W1, M1) than earlier post-operative timepoints (D1), indicating this pre-operative expression was more prognostic of cytokine expression much later in the LT process (Figure 3A).

**Figure 3:**
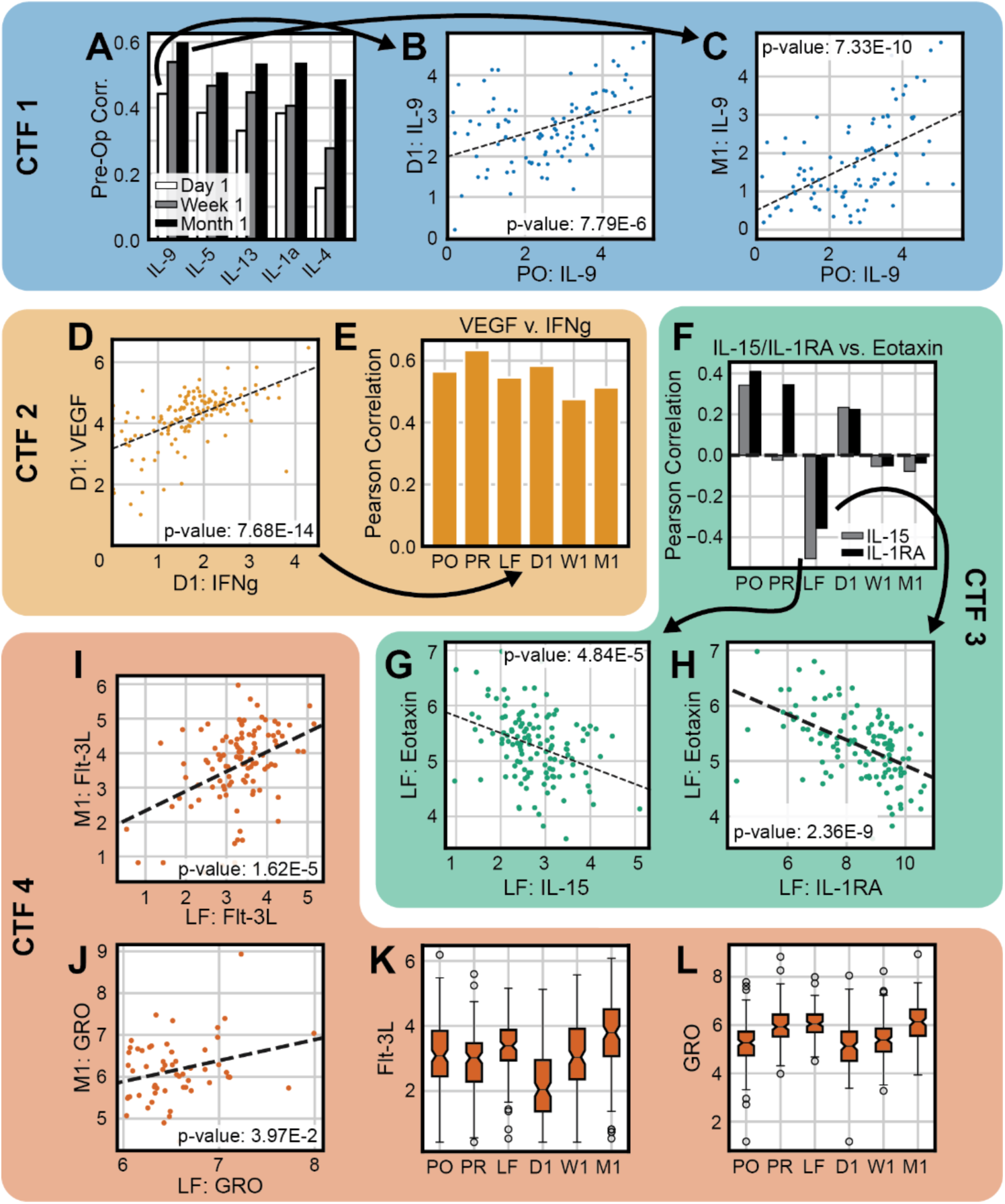
CTF Captures Relationships Between Temporally Disparate LIRI Mechanisms. A) Pearson correlations between LT recipient pre-operative and one-day, one-week, and one-month post-operation measurements for cytokines associated with CTF 1. B–C) Scatter plots comparing IL-9 expression in LT recipients pre-operatively and at one-day post-operation (B) and one-month post-operation (C). D) Scatter plot comparing LT recipient IFNγ and VEGF expression at one-day post-operation. E) Pearson correlations between VEGF and IFNγ expression. F) Pearson correlations between IL-15/IL-1RA and Eotaxin expression. G–H) Scatter plots comparing IL-15 (G) and IL-1RA (H) expression against Eotaxin expression at liver flush. I–J) Scatter plots comparing liver flush and one-month post-operation measurements for Flt-3L (I) and GRO (J) expression. K–L) Boxplots depicting FLT-3L (K) and GRO (L) expression at each measurement timepoint. Dashed lines in scatter plots correspond to the best-fit linear regression line; *p*-values were derived via Pearson correlation.

Finally, to identify if CTF 1’s immunological signatures were driven by etiology or resulted in a specific type of LIRI, we evaluated correlations between patient associations to CTF 1 with different clinical characteristics collected as part of the LT process; collected clinical variables are shown in Table T1. CTF 1 was negatively associated with donor age and donor risk index^28^ (Figure 4A–B), indicating this recurring cytokine signature manifested primarily in LTs with younger, low-risk donor livers.

**Figure 4:**
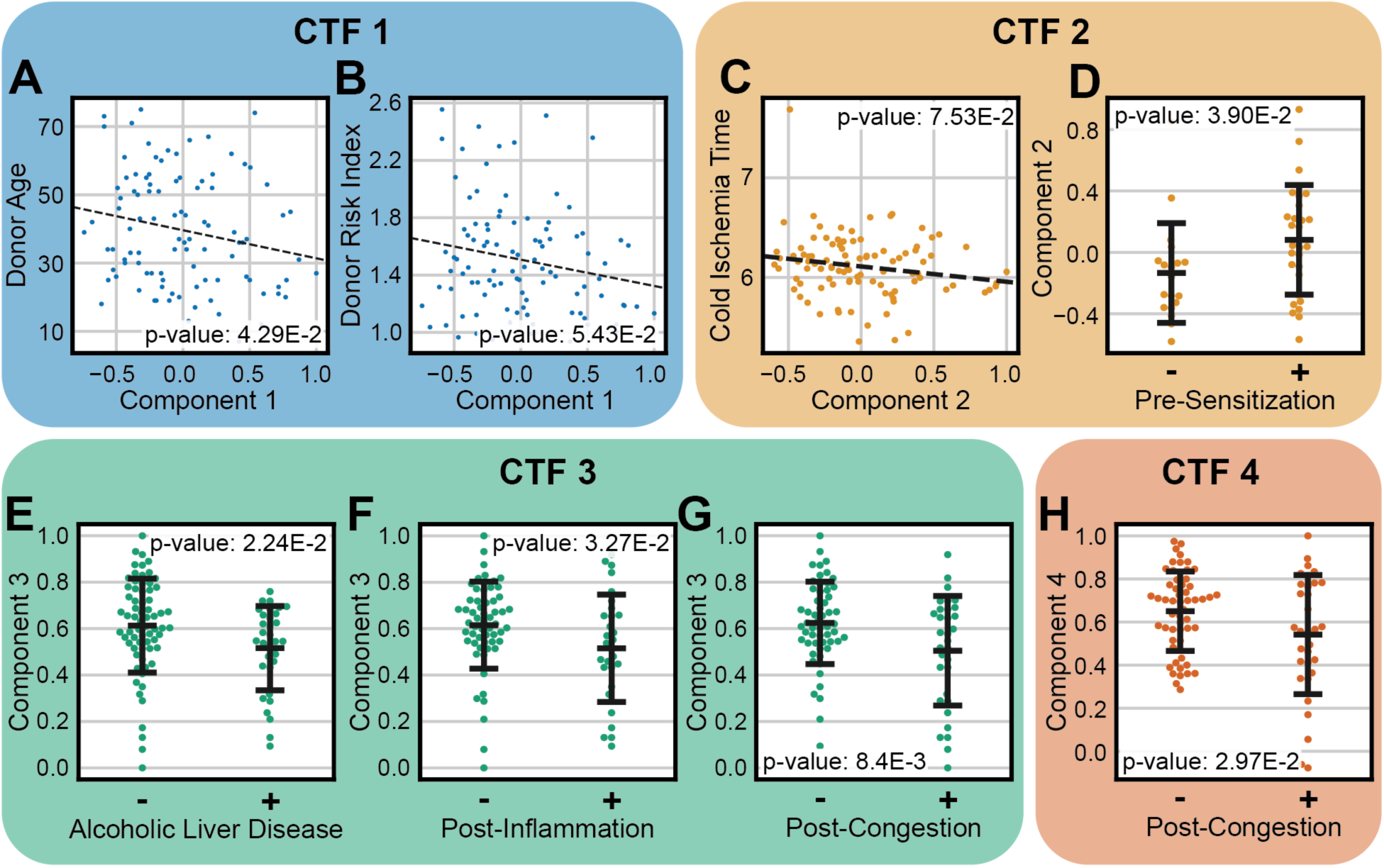
CTF Links LIRI Mechanisms to Clinical Phenotypes. A–B) Scatter plots comparing patient associations to CTF 1 with donor age (A) and donor risk index (B). C) Scatter plot comparing patient associations to CTF 2 with cold ischemia time. D) Swarm plot comparing patient associations to CTF 2 between patients with and without pre-sensitization. E-G) Swarm plots comparing patient associations to CTF 3 between patients with and without alcoholic liver disease (E), high post-transplant inflammation (F), and high post-transplant sinusoidal congestion (G). H) Swarm plot comparing patient associations to CTF 4 between patients with and without high post-transplant sinusoidal congestion. Dashed lines in scatter plots correspond to the best-fit linear regression line. For scatter plots, correlations between patient component associations and clinical variables were derived via Pearson correlation. For swarm plots, average component associations were compared between patient subsets via independent *t*-test.

**Table T1:**
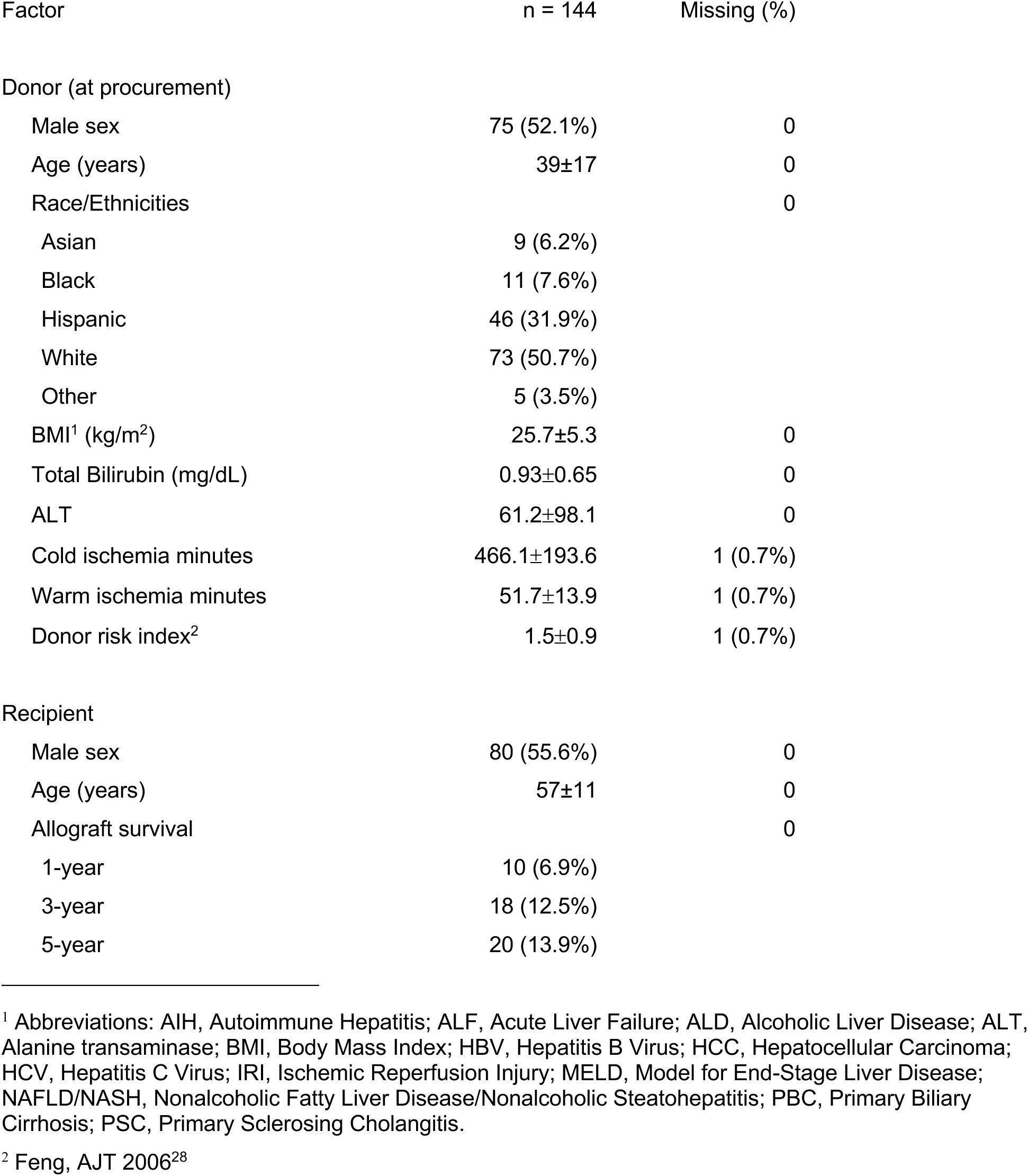
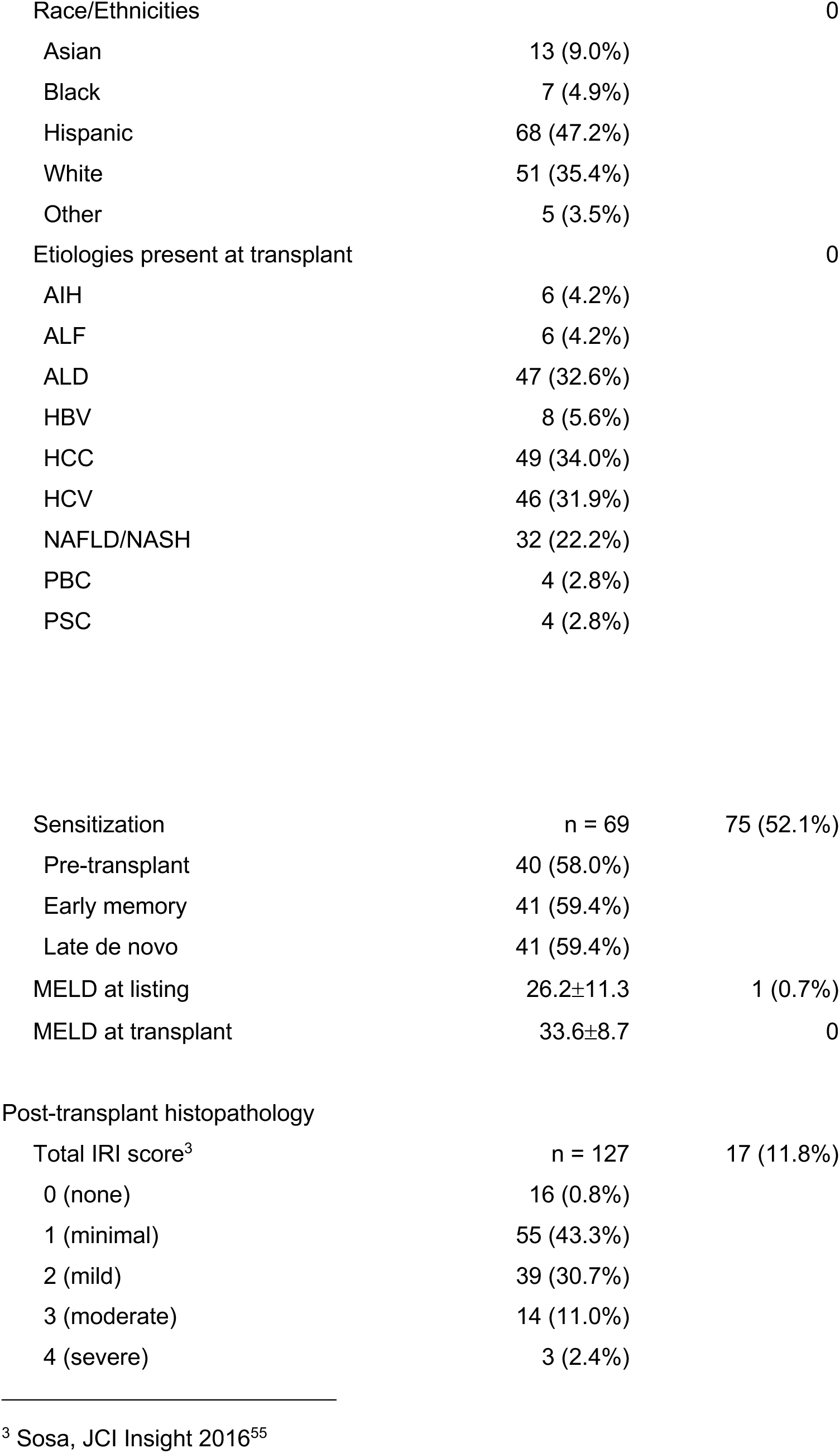
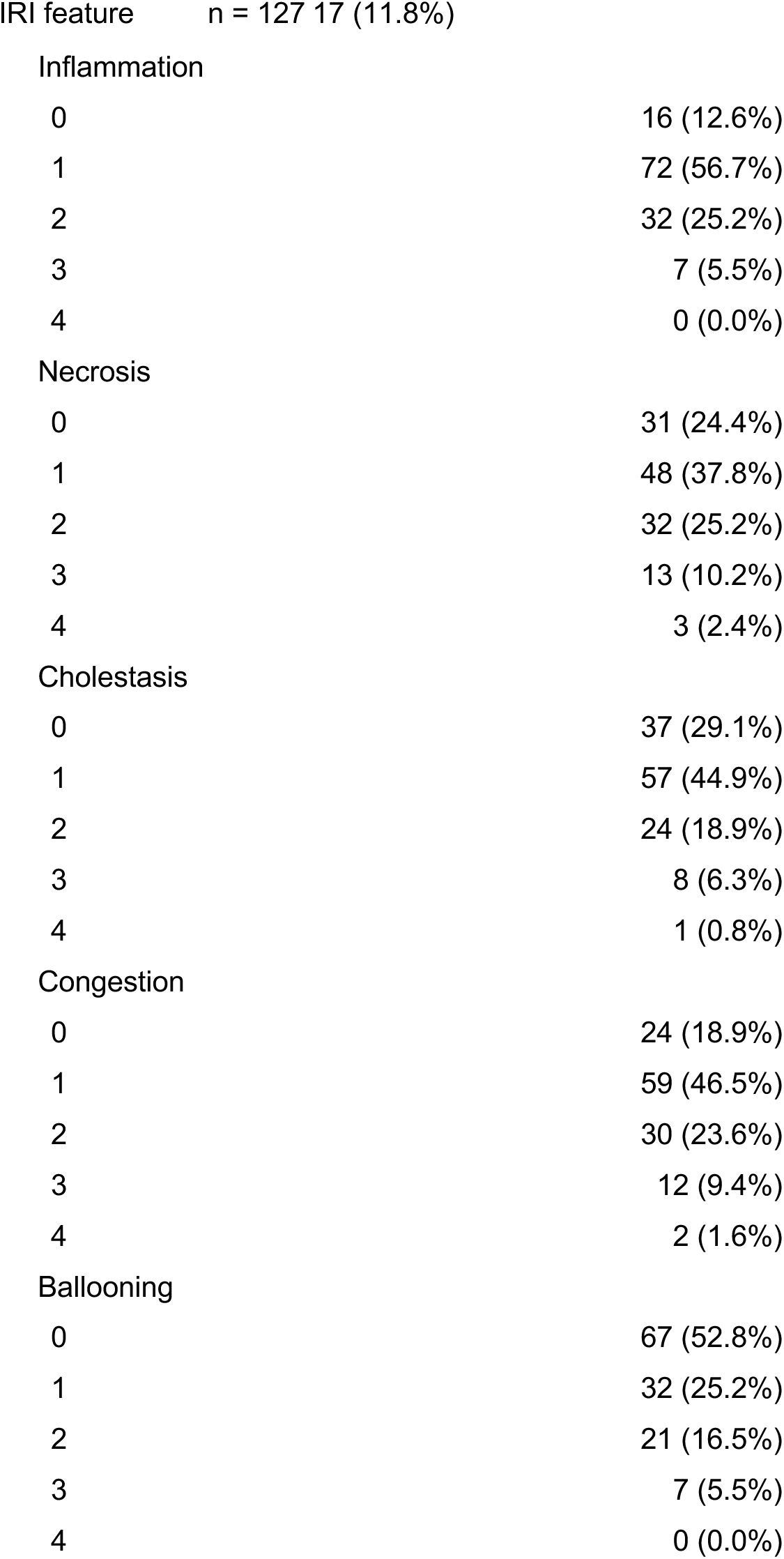
Clinical Characteristics of Liver Transplantation Study Population. Values are arithmetic means ± standard deviation for continuous variables or frequencies (%) for categorical variables.

Comparatively, CTF 2 associated with elevated VEGF, IL-12P40, IFNγ, IL-17A, IL-12P70, and IL-7 early in LT that moderately attenuated one-day post-operation (Figure 2E, G). This signature also negatively associated with many of the cytokines positively associated with CTF 1 (IL-1a, IL-4, IL-5, IL-9, IL-13), indicating that CTF 2 may involve signatures regulatory of those in CTF 1. Collectively, these cytokine signatures corresponded to elevated ALT over the first post-operative week (Figure 2F, H). Clinically, CTF 2 was both negatively associated with cold ischemia time and significantly greater for LTs with pre-sensitized recipients that exhibited human leukocyte antigen (HLA) antibodies, indicating that this immunological signature was (1) more pronounced in LT recipients pre-inclined to alloimmune responses and (2) muted in LTs with extended cold ischemia (Figure 4C-D). Furthermore, in combining IFNγ and VEGF into one component, CTF 2 indicated expression of the two cytokines may be linked. Comparing IFNγ and VEGF expression across patients found that IFNγ and VEGF expression were tightly correlated across timepoints, showing a potential link between the two cytokines in response to LT (Figures 3D-E).

CTF 3, conversely, highlighted a transient signature that manifested primarily at the time-of-transplant. This component associated with elevated Eotaxin and reduced IL-1RA and IL-15 at the LF and D1 timepoints (Figure 2E, G) that corresponded to elevated TBIL scores over the first post-operative week (Figure 2F, H). Comparing Eotaxin expression with IL-15 and IL-1RA indeed found that Eotaxin expression was negatively associated with IL-1RA and IL-15 expression exclusively at the LF timepoint, indicating that elevated Eotaxin shortly after transplant associated with impeded IL-1RA and IL-15 expression (Figure 3F-H). Patients without high post-transplant inflammation or sinusoidal congestion had significantly greater CTF 3 associations (Figure 4F–G), indicating this Eotaxin-based response associated with reduced forms of some post-transplant LIRI. This signature was also lower for LT recipients with alcoholic liver disease (ALD) (Figure 4E).

CTF 4 associated with elevated TBIL in the first week post-operation, like CTF 3, but reflected a unique cytokine signature (Figure 2E–H). This component associated with Flt-3L and GRO that were elevated at the LF and M1 timepoints but reduced at the PO and PR timepoints (Figure 2E, G). Like CTF 1, CTF 4’s associations to early and late measurement timepoints indicated a recurring cytokine signature. In examining the raw cytokine measurements, Flt-3L and GRO both exhibited elevation in expression at LF and M1 with a noticeable dip at the D1 timepoint (Figure 3K–L). Furthermore, comparing Flt-3L and GRO expression at LF and M1 confirmed that expression at M1 was strongly correlated to expression at LF, suggesting that early elevation in Flt-3L and GRO predisposed for elevated Flt-3L and GRO much later post-operation (Figure 3I–J). Like CTF 3, this signature was more pronounced in patients without post-transplant sinusoidal congestion (Figure 4H).

With the CTF components revealing correlations between immunological signatures and clinical variables, we next evaluated correlations between raw cytokine expression and clinical variables to identify if these clinical correlations are found only through CTF integration (Figures S4–S5). Some CTF clinical associations—such as the negative correlation between CTF 1 (IL-4, IL-5, IL-13, MCP-3, and TNFβ elevation at PO, D1, and M1) with donor age (Figure 4A)—were also found in correlating individual cytokines against the clinical variables (Figure S4/S5); a benefit of CTF, however, is that the component links these cytokines to a common immunological signature, indicating that the cytokines in CTF 1 share an underlying mechanism negatively correlated with donor age whereas independent statistical tests do not identify potential relationships between cytokines. Furthermore, some clinical correlations—such as the increased associations in CTF 2 (chronic IFNγ, IL-12, and VEGF elevation) for transplants with pre-sensitized recipient (Figure 4D)—were not found in examining individual cytokines (Figure S4/S5), indicating that this integration through CTF more effectively captured immunological signatures associated with etiology and risk factors. Similarly, in evaluating associations between clinical variables (Figure S6), CTF more effectively captured relationships between etiology and LIRI. CTF 3’s linking of ALD with post-transplant sinusoidal congestion and inflammation was not found through clinical correlations, indicating this signature relies upon integration of longitudinal molecular measurements.

### Outcome-Informed Tensor Partial Least Squares Regression (tPLS) Improves the Accuracy of LT Prognosis

With CTF effectively identifying immunological signatures across LT, we next sought to evaluate if these signatures were predictive of LT outcome. In pairing prediction models with the CTF components to predict LT allograft loss, however, we found that the CTF components were not predictive of LT outcome (Figure S7). We hypothesized this limited predictive ability may be a by-product of CTF’s factorization rather than a limitation of the molecular measurements. CTF produces components that most effectively capture variation in the measurements, and this initial exploration identified novel relationships between temporally disparate immunological signatures, LIRI, and etiology. To best capture signatures prognostic of allograft loss, however, these components could be re-tuned to instead capture patterns relating the longitudinal datasets to allograft loss.

To improve recognition of mechanisms related to allograft loss, we exchanged CTF for a coupled form of tensor partial least squares regression (tPLS)^29^. Like CTF, tPLS is a tensor-based factorization technique that reduces a set of coupled tensors to a series of components that capture patterns within the datasets; unlike CTF, tPLS incorporates knowledge of outcome by regressing the longitudinal datasets against allograft loss, therefore producing components that capture the relationship between the longitudinal datasets and LT outcome (Figure 5A). Here, we regressed the longitudinal datasets against five-year allograft survival.

**Figure 5:**
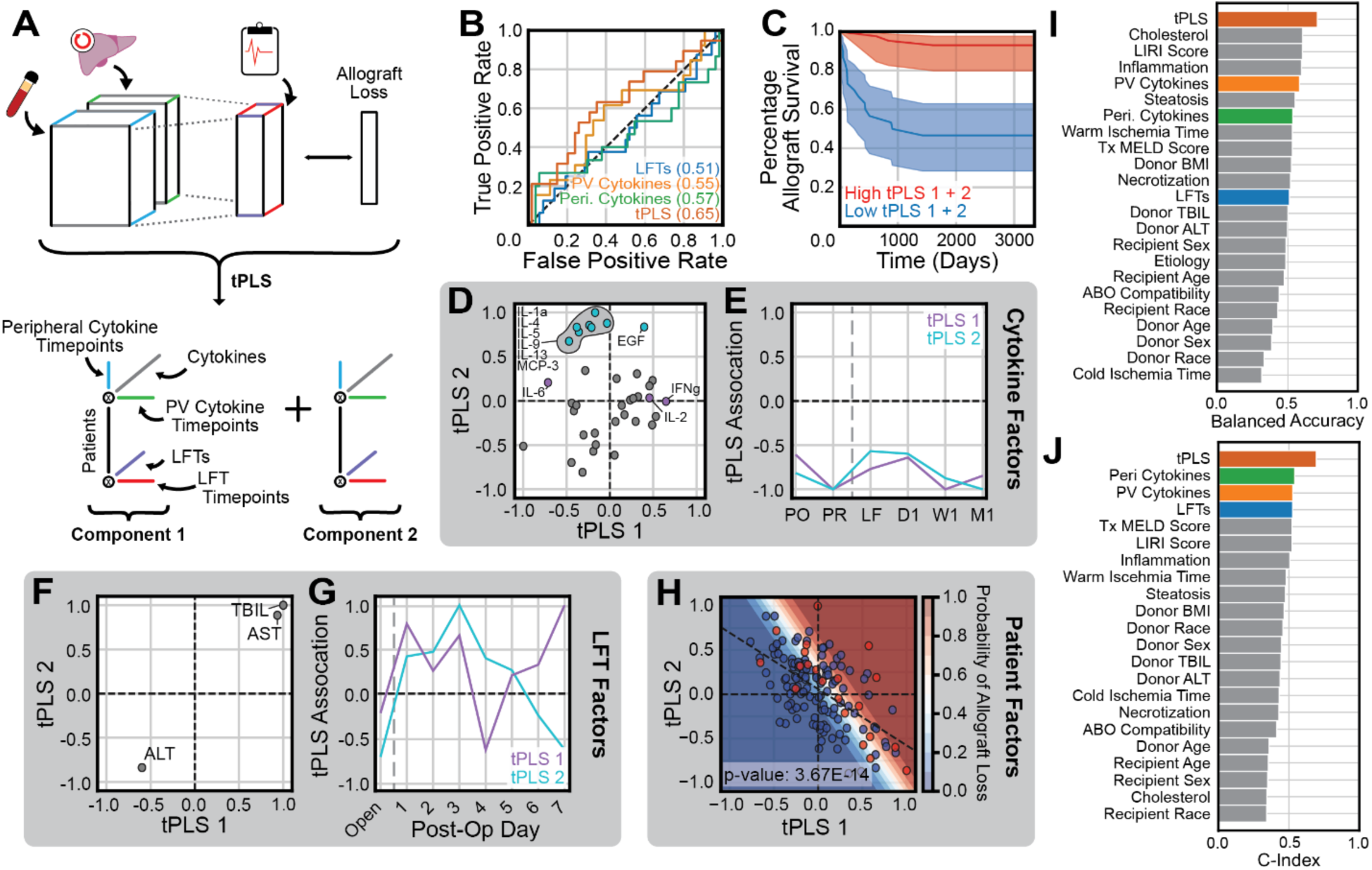
Coupled tPLS Accurately Prognoses 5-Year Mortality. A) Coupled tensor partial least squares (tPLS) jointly regresses the longitudinal datasets against 5-year mortality, producing components that capture the covariance between longitudinal measurements and transplant outcome. B) Receiver-operating characteristic (ROC) curves comparing the coupled tPLS model against tPLS models trained with a single longitudinal dataset. Area under the ROC curve is shown in parentheses. C) Kaplan-Meier curves comparing allograft survival proportions between the 30 patients with the greatest and least associations to tPLS components 1 and 2. Shaded area represents 95% confidence interval for allograft survival proportion. D, F, H) Scatter plots comparing tPLS component associations to cytokines (D), LFT scores (F), and patients (H). The dashed diagonal line in patient scatter plot (H) corresponds to best-fit linear regression line between tPLS components; *p*-value corresponds to Pearson correlation. E, G) Line plots comparing tPLS component associations to cytokine (E) and LFT (G) measurement timepoints. Vertical dashed line corresponds to time-of-transplant. I, J) Bar plots comparing balanced accuracy (I) and concordance index (J) in prognosing 5-year mortality from tPLS, single-dataset, and clinical models.

tPLS requires that the user must specify (1) the number of components to use in factorization and (2) variance scaling between datasets. We chose to optimize these parameters for allograft loss prediction accuracy, resulting in a 2-component tPLS model with increased emphasis (variance multiplied by 100) in peripheral cytokine measurements and decreased emphasis (variance divided by 100) in LFT measurements (Figure S8). The resulting tPLS model predicted allograft loss with a balanced accuracy of 71%, outperforming tPLS models trained with single datasets (Figure 5B, I) and logistic regression models trained with clinical variables often correlated with LT outcome (Figure 5I). The tPLS components also more effectively prognosed allograft survival times than these single dataset and clinical models (Figure 5J), and patients with strong associations to the tPLS components demonstrated significantly lower proportions of allograft survival (Figure 5C).

To interpret the mechanisms captured by these tPLS components, we compared the cytokine (Figure 5D), cytokine timepoint (Figure 5E), LFT score (Figure 5F), LFT timepoint (Figure 5G), and patient (Figure 5H) factors between tPLS components 1 and 2. Higher associations to either tPLS component associated with increased risk of allograft loss, indicating their corresponding immunological signatures both associated with elevated allograft loss risk (Figure 5H). Patient associations to these tPLS components were also negatively correlated, indicating these components may correspond to opposing processes (Figure 5H). tPLS 1 associated with elevated IL-6 alongside reduced IFNγ and IL-2 expression across the LT process (Figure 5D–E). This component also associated with elevated TBIL that heightened at the end of the first post-operative week (Figure 5F–G). Comparatively, tPLS 2 associated with reduced EGF, IL-1a, IL-4, IL-5, IL-9, IL-13, MCP-3, and TNFβ expression pre-operatively and at W1/M1 post-operation.

This component also associated with elevated TBIL that resolved towards the end of the first post-operative week. Notably, tPLS 2 demonstrated similar cytokine associations to CTF 1, but was predictive of 5-year allograft survival while CTF 1 was not. This may be attributed to the limited predictive ability of the remaining CTF components (Figure S7), and that the interactions between tPLS 1 and 2 may be most predictive of 5-year mortality.

To evaluate if these datasets were predictive of mortality earlier post-operation, we regressed the longitudinal molecular measurements against 1- and 3-year mortality outcomes (Figure S9). tPLS more accurately predicts mortality at 1 and 3 years than clinical and single-dataset models, producing a balanced accuracy of 80% and 70%, respectively (Figure S9A, C). Lower area-under the receiver-operating characteristic curve, however, indicated these models may be limited by the smaller percentage of allograft loss cases at these timepoints (Figure S9B, D). As such, we proceeded with 5-year mortality as the regression target to include more allograft loss cases and characterize immunological signatures involved in longer-term allograft survival.

### Allograft Loss Cases Exhibit Sustained Reduction in IFNγ, IL-4, and Regenerative Factors

To better characterize the immunological signatures associated with 5-year allograft loss found by the tPLS model, we compared cytokine expression patterns between the 30 patients with the greatest and least associations to tPLS components 1 and 2. Both components associated with increasing allograft loss risk, indicating both corresponded to processes that increased allograft loss risk. Patients strongly associated with tPLS component 1 exhibited elevated IL-1RA throughout the entire transplant process that was especially pronounced pre-operatively at PO and PR and long post-operation at W1 and M1 (Figure 6A). These LT recipients with elevated IL-1RA also demonstrated universally elevated IL-6 expression (Figure 6B) alongside reduced post-operative IL-2 and IFNγ expression that was especially diminished at D1 (Figure 6B–D). Patients with strong associations to tPLS 2, conversely, demonstrated longitudinal suppression of IL-4, EGF, MCP-3, and TNFβ (Figure 6E-H). This reduction in expression was most notable pre-operatively and at one-month post-operation. Further examination between IL-4, EGF, MCP-3, and TNFβ found IL-4 expression to be tightly correlated with expression of the other three cytokines (Figure 6I-K) at the first post-operative month.

**Figure 6:**
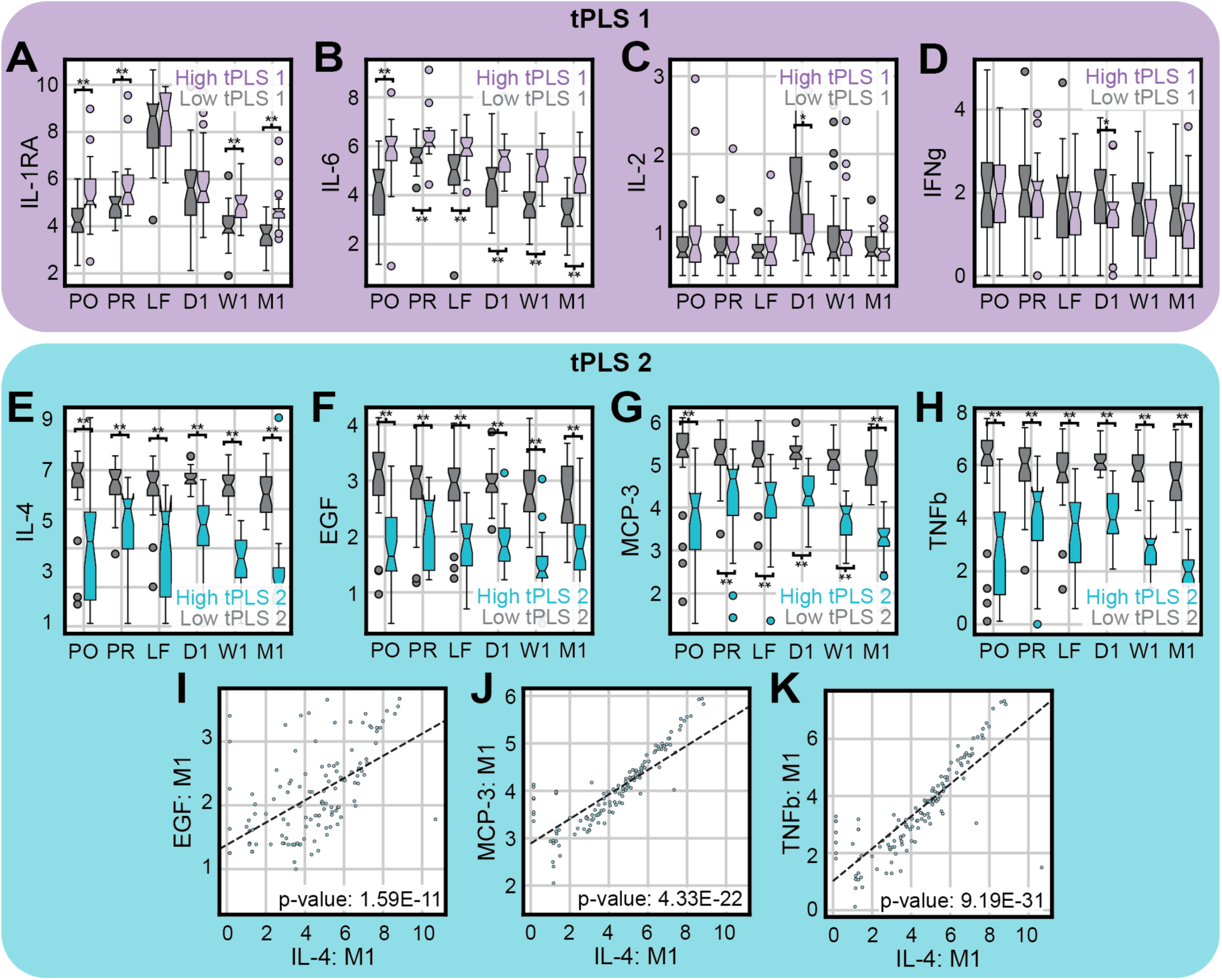
LTs With Allograft loss Exhibit Long-Term Suppression of IFNγ or Th2 Responses. A–D) Boxplots comparing expression of IL-1RA (A), IL-6 (B), IL-2 (C), and IFNγ (D) between the 30 patients with the greatest and least association for tPLS component 1. E–H) Boxplots comparing IL-4 (E), EGF (F), MCP-3 (G), and TNFβ (H) expression between the 30 patients with the greatest and least association for tPLS component 2. I-K) Scatter plots comparing IL-4 expression against EGF (I), MCP-3 (J), and TNFβ (K) expression at the first post-operative month. Dashed lines in scatter plots correspond to best-fit linear regression line. *p*-values in scatter plots were derived via Pearson correlation. Box-plot means were compared via independent *t*-test. * denotes *p*<0.05, ** denotes *p*<0.01 level.

Patient tPLS associations were compared against clinical variables available in the patient meta-data to identify clinical drivers associated with the tPLS components. tPLS 1 was decreased for LT recipients with multiple etiologies (Figure 7A) and hepatocellular carcinoma (Figure 7B) and demonstrated a negative correlation to both donor and recipient age (Figure 7C-D), indicating this signature manifested primarily in LTs featuring both younger donors and recipients. Conversely, tPLS 2 was elevated for LT recipients with non-alcoholic steatohepatitis (NASH) (Figure 7E) and demonstrated significant correlations with both donor age and risk index (Figure 7F-G), indicating this immunological pattern arose primarily in transplants involving marginal livers or livers harvested from older donors.

**Figure 7:**
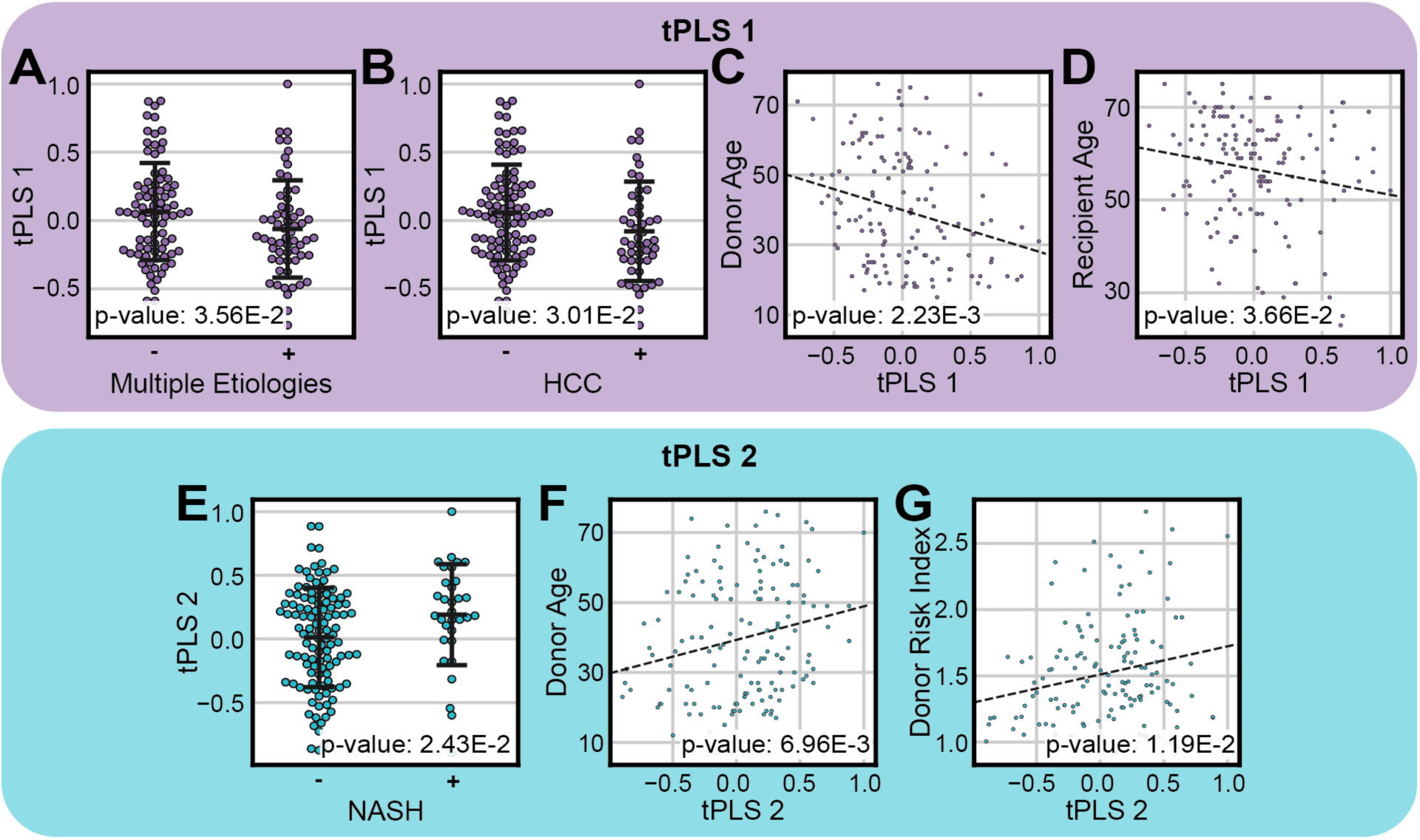
tPLS Immune Signatures Vary with Etiology, Age. A-B) Swarm plots comparing patient associations to tPLS 1 between patients with and without multiple etiologies (A) and hepatocellular carcinoma (HCC) (B). C-D) Scatter plots comparing patient associations to tPLS 1 with donor age (C) and recipient age (D). E) Swarm plot comparing patient associations to tPLS 2 between patients with and without non-alcoholic steatohepatitis (NASH). F-G) Scatter plots comparing patient associations to tPLS 2 with donor age (F) and donor risk index (G). Dashed lines in scatter plots correspond to best-fit linear regression line. *p*-values in scatter plots were derived via Pearson correlation. For swarm plots, average component associations were compared between patient subsets via independent t-test.

## Discussion

While many sources of LIRI have been identified, the relationships between temporally disparate LIRI mechanisms and immunological signatures remain poorly understood, precluding recognition of determinants of liver health and outcome in LT. Here, through tensor-based integration, we identified immunological signatures observed across longitudinal donor and recipient measurements, enabling recognition of relationships between LIRI, immune responses, and disease etiology in LT and their collective associations to eventual transplant outcome. Furthermore, we found that recognition of these signatures is unique to these tensor-based integration approaches as single-dataset and clinical models failed to capture these relationships between immunological pathways, clinical correlates, and later transplant outcome as effectively as these tensor-based approaches.

Some of these signatures corresponded to transient immunological signatures. CTF 3’s strong positive association to Eotaxin—a potent eosinophil chemoattractant^30^—alongside reduced anti-inflammatory IL-1RA at the LF and D1 timepoints may correspond to elevated eosinophil responses shortly post-transplant. The role of such eosinophil responses in LT outcome is still uncertain^31–33^; the present study finds that, while such eosinophil signatures were not associated with allograft survival, they may be protective against some sources of LIRI. This signature was reduced in recipients with alcoholic liver disease (ALD) alongside patients with post-transplant sinusoidal congestion and inflammation, indicating ALD may impede eosinophil responses that were protective against post-transplant LIRI.

Tensor-based integration also found recurring immunological signatures primed pre-operatively that returned post-operatively. We found that elevated expression of GRO and Flt-3L in donor livers immediately post-operation corresponded to elevated GRO and Flt-3L at one-month post-operation, indicating a connection between these signatures at the time-of-transplant with those up to a month post-operatively. Similarly, we found that pre-operative elevation of Th2 cytokines in recipients—such as IL-4, IL-5, IL-9, and IL-13^34^—corresponded to returning Th2 cytokine elevation later post-operation, indicating such pre-operative priming of Th2 responses corresponded to later elevation in Th2 signatures. Curiously, both groups of cytokines exhibited reduced expression post-operation at D1 that returned to elevated levels at W1/M1. Plausibly, this reduced expression could be the result of immunosuppressive therapies. Immunosuppressants are introduced shortly after LT to mitigate risk of early allograft rejection^35^. Many of these immunosuppressants target T cells via IL-2 and can impair both CD4+ Th1 and Th2 responses alongside downstream effector cells that produce GRO and Flt-3L^36,37^.

Regardless of immunosuppression, this study highlights the importance of CD4+ balance in LT. Previous studies have found that pre-operative Th2 elevation associated with reduced risk of acute and chronic LT rejection^34,38,39^. Our analyses expanded upon this signature in tPLS 2, finding that pre-operative suppression of Th2 cytokines—like IL-4, IL-5, IL-9, and IL-13^34^—led to long-term Th2 suppression, elevated allograft loss risk, and reduced expression of growth factors like EGF and TNFβ that are critical in hepatic regeneration^40–44^. Suppression of Th1 responses, however, also associated with increased risk of allograft loss. Patients characterized by tPLS 1 exhibited elevated IL-1RA alongside reduced expression of CD4+ Th1 cell products (IL-2 and IFNγ), indicating that tPLS 1 may correspond to reduced Th1 activity^45,46^. Reductions in Th1 responses associated negatively with Th2 suppression, indicating suppression of either CD4+ archetype led to elevations in the other. Excessive suppression of either archetype, however, associated with increased allograft loss risk, indicating that Th1 and Th2 responses must be balanced for optimal five-year allograft survival. We also observed diverging clinical associations with CD4+ polarization, finding that LTs with younger donors and recipients exhibited reduced Th1 responses while LTs with older and higher-risk donors exhibited reduced Th2 responses, indicating that age and donor liver health may modulate later balances in CD4+ polarization.

These findings also provided further insights into IFNγ’s role in LT. As a pro-inflammatory Th1 cytokine, IFNγ can exacerbate LIRI and increase acute LT rejection risk^17,47–49^. In congruence, we observed that elevated IFNγ associated with elevations in ALT over the first post-operative week. As suppression of IFNγ and Th1 responses associated with allograft loss, however, such inflammatory responses appear to be a critical part of allograft survival. Here, we found that IFNγ expression was tightly correlated with VEGF, a growth and angiogenic factor that supports liver regeneration and angiogenesis^50,51^, therefore indicating IFNγ may be protective against allograft loss via VEGF expression. We also observed that suppressed IFNγ expression corresponded to elevated IL-6, a pleiotropic and often pro-inflammatory cytokine that has been associated with increased risk of acute and chronic rejection in LT^52,53^.

Furthermore, clinical correlates provided actionable insights into such IFNγ/Th1 immunotolerance mechanisms and the relationships between etiology and later immunological signatures. IFNγ/VEGF signatures were elevated in patients with pre-sensitization—those that exhibited some donor-specific antibody (DSA) response prior to transplant—and somewhat reduced by cold ischemia, indicating pre-sensitizing LT recipients and minimizing cold ischemia time were essential to preserving IFNγ responses. Conversely, Th2 responses appeared more dependent on etiology as Th2 suppression was more pronounced in LT recipients with non-alcoholic steatohepatitis (NASH). Likewise, LT recipients with alcoholic liver disease (ALD) exhibited reduced Eotaxin and elevated post-transplant LIRI, highlighting that deviant eosinophil responses may be a component of post-transplant LIRI for patients with ALD. These etiological correlates, as well as the prior clinical correlates, were found only through tensor-based integration of longitudinal LFT and cytokine measurements, indicating integration of these measurements improved our ability to capture such critical clinical manifestations.

We acknowledge that this study is not without its limitations. First, this study consists of a moderately sized sample of 144 LTs performed at a single center. Thus, these findings outline trends in LT observed at one location for further validation in other geographically independent cohorts. Second, our transplants were missing some measurements across the coupled longitudinal datasets, and some were missing entire timepoints of measurements. While coupled tensor methods can capture patterns in datasets with missing measurements, the absence of some measurements may have precluded recognition of immunological patterns in LT and allograft outcome; notably, the PV cytokine measurements 1) demonstrated the highest amount of missingness and 2) were less-weighted in predicting allograft loss via tPLS than other measurement types. Thus, further investigation is warranted into outcome-associated patterns observed in PV cytokine measurements. Third, as treatment regimens were unavailable to the investigators, the effect of therapeutics and treatment time-courses on these findings is unclear. The immunosuppressive regimens often prescribed to LT recipients, along with their withdrawal, are known to modulate immunological signatures; for example, withdrawal from LT immunosuppressants typically elicits NK cell expansion^54^. Thus, it is unknown if these findings may reflect variation in treatment regimens and further investigation is necessary to determine how therapeutics may interact with the immunological signatures in the present study. Similarly, further investigation is warranted to determine if these patterns are specific to LT or may be generalizable across other transplants involving similar immunosuppressive treatments.

Collectively, these results find that tensor-based integration of longitudinal clinical and cytokine measurements can improve our understanding of the relationships between injury patterns and eventual LT outcome. CTF integration of longitudinal measurements collected across the LT process captured four immunological patterns observed in LTs. Through interpretation of these patterns, we identified relationships between temporally disparate immunological signatures and etiological/injury-driven damage mechanisms, enabling us to recognize relationships between early and later sources of LIRI that were not found via single-timepoint and single-dataset models. Incorporation of outcome through tPLS further reduced these four components to two strongly associated with five-year mortality that similarly out-performed single-dataset and clinical models in LT prognosis. Through examination of these two tPLS components, we identified the importance of balanced Th1 and Th2 responses in preventing LT allograft loss. In combining the tPLS and CTF results, this work also provides further context to the role of Th2 responses and IFNγ in LT outcomes. These efforts demonstrate the importance of profiling and integrating longitudinal clinical and molecular measurements in LT, finding that these tensor-based integration methods enable a more comprehensive overview of injuries accrued over the LT process and their relationship to eventual transplant outcome.

## Methods

### Patients and sample collection

#### Sample Collection

This study consisted of 144 liver transplantation (LT) recipients recruited at UCLA Health between 2013 and 2020. Routine care and immunosuppressive therapies were provided in accordance with UCLA LT protocols. All patient samples were deidentified before use. Patient samples were processed within 4-8 hours of collection and stored at -80⁰C until first use.

As part of standard care, liver function tests (LFTs)—namely, bilirubin (TBIL), alanine transaminase (ALT), and aspartate transaminase (AST)—were collected pre-operation and daily for the first post-operative week from recipient peripheral venous blood. Recipient venous blood was also collected using acid-citrate-dextrose anticoagulant pre-operation (PO) and at one-day (D1), one-week (W1), and one-month (M1) post-operation. Intraoperative blood was collected from the recipient portal vein (PV) prior to reperfusion (PR) and from the donor liver PV as it was flushed through the vena cava during reperfusion (liver flush, LF).

Donor livers were harvested at the time of brain death or at the time of circulatory death via standard techniques. Protocol Tru-Cut needle biopsies were collected from the left lobe intraoperatively after allograft revascularization approximately two hours after reperfusion but before surgical closing of the recipient abdomen.

#### Cytokine Measurements

Plasma samples were prepared via centrifugation at 500 g for 12 minutes and stored at 80⁰C. Human 38-plex cytokine kits (EMD Millipore, HCYTMAG-60K-PX38) were used per manufacturer instructions to assay cytokine and chemokine expression in collected blood samples. The complete panel screened for expression of the following cytokines: chemokine (C-X-C motif) ligand 1 (GRO); Eotaxin; epidermal growth factor (EGF); fibroblast growth factor 2 (FGF-2); FMS-like tyrosine kinase 3 ligand (Flt-3L); Fractalkine; granulocyte colony stimulating factor (G-CSF); granulocyte-macrophage colony stimulating factor (GM-CSF); Interferon alpha 2 (IFNα2); Interferon gamma (IFNγ); Interferon gamma-induced protein 10 (IP-10); Interleukins 1a, 1b, 1RA, 2, 4, 5, 6, 7, 8, 9, 10, 12P40, 12P70, 13, 15, and 17A (IL-1a, IL-1b, IL-1RA, IL-2, IL-4, IL-5, IL-6, IL-7, IL-8, IL-9, IL-10, IL-12P40, IL-12P70, IL-13, IL-15, and IL-17A); macrophage-derived chemokine (MDC); macrophage inflammatory protein 1 beta (MIP-1β); monocyte chemoattract proteins 1 and 3 (MCP-1 and MCP-3); soluble cluster of differentiation 40 ligand (sCD40L); transforming growth factor alpha (TGFα); tumor necrosis factors alpha and beta (TNFα and TNFβ); and vascular endothelial growth factor (VEGF). Concentrations were calculated using Milliplex Analyst software version 4.2. Luminex assays and analysis were performed by the UCLA Immune Assessment Core.

#### Histopathological Scoring

Following reperfusion, needle biopsy samples were formalin-fixed, paraffin-embedded and stained with hematoxylin and eosin following a standardized protocol established by the Translational Pathology Core Laboratory at UCLA. Biopsies were reviewed by hepatopathologists and graded semi-quantitatively for inflammation, necrosis, sinusoidal congestion, ballooning, and cholestasis in accordance with our prior scoring criteria^55^. For each biopsy, each type of liver ischemia reperfusion injury (LIRI) was graded on a scale of 0-4. Scores of 0-1 indicate none or minimal of the corresponding LIRI; scores 2-4 indicate a mild or severe case of the corresponding LIRI type. Scores of 2+ were used to define presence of the corresponding LIRI type.

#### Donor Risk Index

To quantify LT allograft loss risk from donor factors, the donor risk index was calculated for each transplant^28^. This index was calculated according to the following equation:

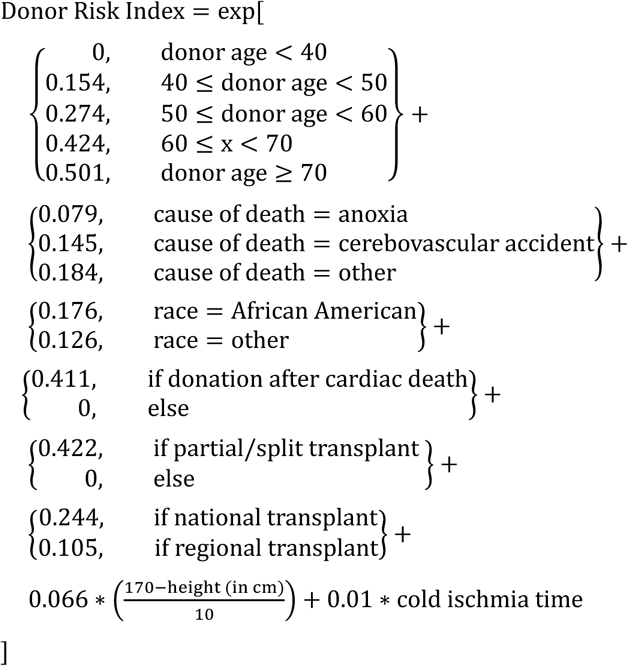

### Computational Methods

#### Measurement Preprocessing

Before factorization, the peripheral blood cytokine, PV cytokine, and LFT measurements were separated into their own matrices. To account for differing ranges of values across measurement types and sources, measurements were power-transformed then mean-centered such that each cytokine/LFT had a mean of zero across all patients and measurement timepoints. To ensure equal variance across measurements, each cytokine/LFT was then divided by its variance to enforce an equal variance of 1 across patients and timepoints. This normalization procedure was performed separately for each dataset.

Following normalization, the variance of each dataset can be adjusted to modify its emphasis in the resulting factorization as coupled factorization methods will over-emphasize datasets with larger variance in the resulting factor matrices. For coupled tensor factorization (CTF), we enforced equal variance across datasets to ensure each dataset tensor (peripheral cytokines, PV cytokines, and LFTs) was weighted equally in the resulting factorization. For tensor partial least squares (tPLS), we choose these variances to optimize five-year mortality prediction accuracy.

#### Patient Selection for Model Training

To prevent the coupled tensor factorization (CTF) model from potentially emphasizing patterns related to data missingness, patients missing all measurements for at least one timepoint in the cytokine or liver function test (LFT) datasets were removed prior to factorization. This resulted in measurements from 100 transplants used to fit the CTF model.

Given the relatively small proportion of allograft loss cases, transplants with allograft loss and missing measurements were not removed prior to factorization via tensor partial least squares (tPLS) as such rare events were needed to effectively fit the tPLS model. Transplants from non-allograft loss cases missing all measurements for at least one timepoint, however, were removed. Additionally, as patients were admitted to the study on a rolling basis, non-allograft loss cases that were collected within 5 years of the last clinical follow-up were excluded from tPLS model fitting as 5-year survival could not be determined. This resulted in measurements from 73 transplants used to fit the tPLS model; removal of patients from the tPLS model occurred prior to data normalization, model training, and cross-validation. Transplants excluded from the model fitting were later transformed via the fitted tPLS model to derive patient associations to the tPLS components for the excluded transplants. To ensure equal comparisons between tPLS and non-tPLS models that cannot handle missing data, allograft loss accuracy and concordance index metrics only consider these 73 patients.

#### Cross-Validation, Oversampling, and Accuracy Evaluation

Matrix and tPLS model accuracies in predicting allograft loss were evaluated via 10-fold stratified cross-validation, as implemented in scikit-learn^56^. Briefly, 10% of the transplant measurements were withheld as a “testing” dataset while the remaining 90% were used to train the model as a “training” dataset. Stratification ensured that training and testing datasets had equal proportions of allograft loss cases. Following training against the training dataset, the fitted model was then applied to the testing dataset to predict allograft outcomes for the withheld transplants. This process was repeated nine additional times, each time choosing a different testing dataset, such that each transplant received exactly one prediction for allograft outcome. Matrix and tPLS models were re-fit between cross-validation folds to prevent data leakage.

To mitigate the effects of class imbalances in model fitting, oversampling was implemented in the cross-validation schemes for the tPLS and flattened models. In the above cross-validation schemes, after defining testing and training datasets, allograft loss cases in the training datasets were oversampled; allograft loss cases in the training dataset were sampled with replacement until the training dataset had an equal proportion of allograft loss and non-allograft loss cases. This oversampled training dataset was then used to train the model before application to the testing dataset. Testing datasets were never resampled.

#### Paired tPLS Models

Logistic regression (LR) and Cox proportional hazards (CPH) models were used to predict allograft survival and allograft survival times from tPLS patient factors, respectively. LR models were implemented via scikit-learn^56^; CPH models were implemented via lifelines^57^. As specified above, patient associations to tPLS components were derived via stratified cross-validation with oversampling. Within each cross-validation loop, LR and CPH models were fit to the patient factor from the training dataset; the fitted LR and CPH models were then applied to the testing dataset’s patient factor to predict allograft survival and expected allograft survival times for each patient. LR and CPH models were re-fit with tPLS models each cross-validation fold to ensure no data leakage occurred.

#### Matrix Models

To provide a baseline of comparison for the tPLS models, matrix-based classification models were used via LR. Cytokine and LFT measurements were flattened into matrices for these classification models; measurement timepoints and cytokines (for the peripheral and PV cytokine measurements) or LFT scores (LFT measurements) were concatenated into one dimension, producing a matrix with rows corresponding to transplants and columns corresponding to the merge of measurements and measurement timepoints. LR was applied to this matrix to predict allograft loss via cross-validation and oversampling, as outlined above.

To compare abilities in prognosing allograft survival times, CPH models were implemented. Tensor-structured datasets were flattened as described above and then used to train and predict expected allograft survival from molecular measurements via the previously described cross-validation and oversampling schemes.

#### Reconstruction Fidelity (R2X)

To evaluate the ability of CTF to effectively reconstruct the datasets, we calculated the variance explained by CTF in R2X. First, total variance was derived as the sum of squared Frobenius norms across datasets: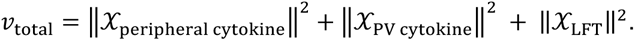 We then derived the squared Frobenius norms of the difference between the tensor datasets and their reconstructed forms: 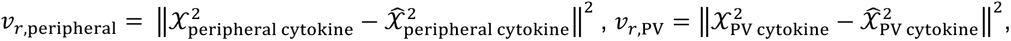 and 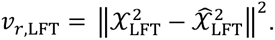 The variance explained is then calculated as 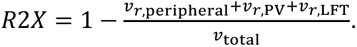 Missing values were ignored in the above calculations.

#### Balanced Accuracy

As the datasets had an over-representation of non-allograft loss cases, balanced accuracy was used instead of accuracy to more effectively manage class imbalances and evaluate model performance. To calculate balanced accuracy, we derived the recall for both the allograft loss and non-allograft-loss cases; balanced accuracy is the arithmetic mean of recall scores across allograft and non-allograft-loss cases.

#### Factor Match Score (FMS)

To evaluate the consistency of the CTF factorization and its captured patterns across LTs, we calculated the factor match score (FMS) for CTF using the Python package TLViz^58^. First, we derived the factor matrices for CTF applied against all measurements. We then sampled LTs with replacement, resulting in a resampled dataset of equal size to the original CTF dataset with a different composition of patients. CTF was then applied to this resampled dataset to produce a second set of factor matrices.

To evaluate the consistency in factorization across the cytokine and LFT measurements, the FMS was calculated separately for cytokine and LFT measurements. CTF components are not sorted in order of variance explained, and the order of components can vary by decomposition; two decompositions may produce the same components but in different order. To address this permutation dependency, linear sum assignment via SciPy was used to re-order components in the resampled CTF factor matrices to best match the components between factor matrices. Following component reordering of the resampled factor matrices, the dot product of the default and resampled CTF components was divided by the product of the component Frobenius norms for each factor matrix, then multiplied across factor matrices and summed across components. FMS is calculated separately for LFT and cytokine factors:

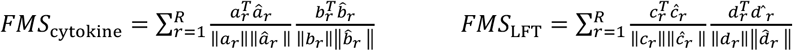

where a, b, c, and d denote the component vectors with regards to cytokine, cytokine timepoint, LFT score, and LFT timepoint, respectively. ^ denotes resampled CTF vector.

The above results in separate FMS scores for cytokine and LFT factors that range between 0 (completely orthogonal) and 1 (identical components). As the resampled dataset consists of a different make-up of LTs, the patient dimension is ignored in these calculations.

#### Statistical Methods

Independent *t*-tests, Pearson correlations, and Fisher’s Exact tests were implemented via SciPy^59^. Two-sided independent *t*-tests were used to compare means of component associations and cytokine expression across subsets of patients. Pearson correlation was used to compare correlations between continuous values. Fisher’s Exact test was used to evaluate association between binary clinical variables.

#### Coupled Tensor Factorization

We implemented Coupled Tensor Factorization (CTF) to jointly analyze the three longitudinal datasets: the peripheral blood cytokine (PB), portal vein cytokine (PV), and liver function tests (LFT) measurements. Each dataset was organized into a three-mode tensor as *X*_PB_, *X*_PV_, and *X*_LFT_, respectively. *X*_PB_ is organized by patients, cytokines, and PB measurement timepoints; *X*_PV_ is organized by patients, cytokines, and PV measurement timepoints; *X*_LFT_is organized by patients, LFT scores, and LFT measurement timepoints. To simplify computational implementation, *X*_PB_ and *X*_PV_ were coupled across their shared cytokine dimension by concatenating the two three-mode tensors across their shared patient and cytokine modes, leading to a concatenated cytokine tensor *X*_Cytokine_ with dimensions of patients, cytokines, and cytokine timepoints that contains both the PB and PV cytokine timepoints. To discover signals within and across datasets, each tensor was then reduced and decomposed into *R* Kruskal-formatted tensors in a coupled fashion. Specifically, each tensor is approximated as

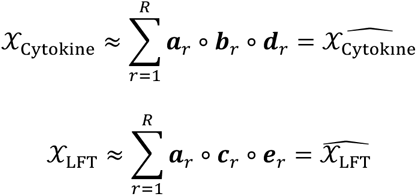

where ***a****_r_* is the *r*-th factor for the patient mode, ***b****_r_* for the cytokine mode, ***c****_r_* for the LFT score mode, ***d****_r_* for the cytokine time mode, ***e****_r_* for the LFT time mode. Each factor is a vector that has the same length as the number of distinct entries in its respective mode, and “∘” represents the vector outer product. Concatenating all *R* factors for each mode, we have the factor matrices *A*, *B*, *C*, *D*, and *E*, respectively. Notice that the patient factors, ***a****_r_*, are shared by both tensors. This is a coupled decomposition, where these tensor decompositions must be solved jointly^27^.

This coupled tensor decomposition was solved through an alternating least squares scheme. Each factor matrix was first initialized by an imputed singular value decomposition (SVD) of the unfolded tensor. For uncoupled modes included in only one tensor—namely, the cytokine (*B*), LFT score (*C*), and the two temporal modes (*D* and *E*)—SVD was performed on the unfolded form of the corresponding tensor. For example, the LFT score mode (*C*) was initialized as

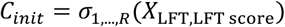

where *σ*_1,…,*R*_ represent the *R* largest singular vectors, and *X*_LFT,LFT score_ is the LFT tensor unfolded into a matrix along the LFT score mode (denoted by mode after the comma). Thus, the uncoupled time factor matrices were initialized as

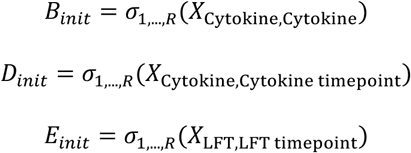

For the coupled patient mode included in every tensor, SVD was performed on a matrix consisting of all tensors unfolded and concatenated into one matrix. Thus, the coupled patient factor matrix was initialized as

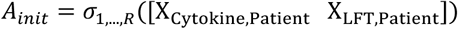

where “[ ]” indicates the column concatenation of the matrices within it.

After initialization, least squares solving was performed for each factor matrix until convergence. For uncoupled modes, factor matrices were updated to the least squares solution of the Khatri-Rao product (indicated by “⊙”) of the other factor matrices in the corresponding tensor and the unfolded data tensor. For example, the LFT factor matrix (*C*) was updated as the least squares solution of the LFT tensor unfolded along the LFT score mode (*X*_LFT,LFT Score_) and the Khatri-Rao product of the patient (*A*) and LFT timepoint (*E*) factor matrices:

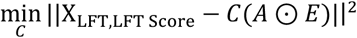

Thus, the other uncoupled factors were updated as:

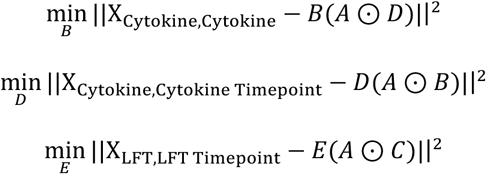

Likewise, the coupled patient mode was updated with the least squares solution of (1) the matrix consisting of all tensors unfolded and concatenated into one matrix and (2) the concatenated matrix of Khatri-Rao products of factor matrices for each tensor:

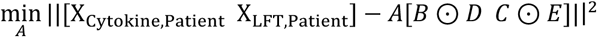

Following each least squares update, each factor matrix was normalized to have an L2-norm of 1. Iterations were repeated, updating each factor, until the variance explained (R2X) improved by less than 1E-8 between iterations. Missing values were ignored throughout. Following convergence, *D* was unstacked to recover component associations to PB and PV cytokine timepoints separately.

### Coupled Tensor Partial Least Squares

To capture patterns between longitudinal datasets and transplant outcome, we implemented a coupled form of tensor partial least squares (tPLS). As with CTF, each longitudinal dataset was organized into a three-mode tensor and the PB and PV cytokine datasets were concatenated into one single cytokine tensor. tPLS then reduced the coupled, tensor-structured longitudinal datasets to a series of *R* Kruskal-formatted tensors:

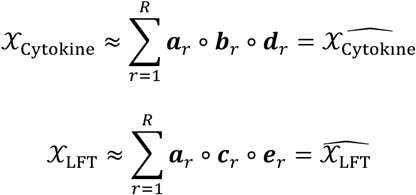

where *X*_Cytokine_ and *X*_LFT_represent the tensor-structured longitudinal cytokine and LFT measurements, respectively, and ***a****_r_* is the *r*-th factor for the patient mode, ***b****_r_* for the cytokines mode, ***c****_r_* for the LFT score mode, ***d****_r_* for the cytokine timepoint mode, and ***e****_r_* for the LFT time mode. Retaining the notation from CTF, concatenating all *R* factors for each mode produces the factor matrices, *A*, *B*, *C*, *D*, and *E*, respectively. Again, the patient factors, ***a***, are shared by both tensors, indicating this is a coupled decomposition where the tensor decompositions must be solved jointly.

In addition to the above factor matrices, knowledge of outcome was incorporated into the tPLS factorization via a vector of binary values, *y*, corresponding to five-year allograft mortality for each patient. Like the measurement tensors, *y* was approximated as the outer product of *R* Kruskal-formatted tensors:

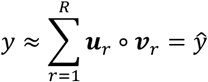

where ***u****_r_* is the *r*-th factor that defines the projection of patients to factors and ***v****_r_* defines the projection between the factors and allograft survival. Similarly, all *R* factors can be concatenated to produce factors matrices *U* and *V*, respectively.

This outcome-informed coupled factorization can be solved using a variant of the nonlinear iterative partial least squares (NIPALS) algorithm adjusted for tensor-structured datasets first described by Bro^29^. Each factor matrix was first initialized with zeros except for *U* where each factor is initialized with *y*.

Solving then proceeds iteratively, resolving each component, *r*, sequentially. First, ***u****_r_* was projected onto each dataset tensor *X*:

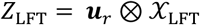

where ⊗ indicates tensor product. *Z*_Cytokine_ and *Z*_LFT_, the projections of *u_r_* onto *X*_Cytokine_ and *X*_LFT_, respectively, were then decomposed via rank-one canonical polyadic decomposition to update ***b***_***r***_, ***c***_***r***_, ***d***_***r***_, and ***e***_***r***_. This decomposition was performed via alternating least squares, iterating between solving for the least squares solutions of regressing *Z*_Cytokine_ and *Z*_LFT_ against ***b***_***r***_/***d***_***r***_ (for *Z*_Cytokine_) and ***c***_***r***_/***e***_***r***_ (for *Z*_LFT_) until convergence was reached:

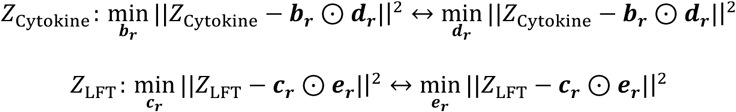

The dataset tensors *X*_Cytokine_ and *X*_LFT_ were then unfolded along the patient dimension. The shared patient factor, ***a****_r_*, was updated to the arithmetic mean of least-squares solutions in regressing each unfolded dataset tensor against its respective factors:

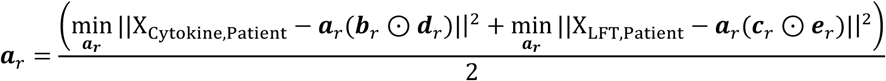

***v****_r_* was updated to the least squares solution of *y* regressed against ***a****_r_*

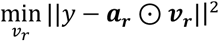

Finally, ***u****_r_* was updated to the least squares solution of *y* regressed against ***v****_r_*

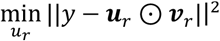

Following each update, each factor was normalized to have an L2 norm of 1. Iterations were repeated until convergence, or the norm of the difference between factors in consecutive iterations was less than 1E-8. Missing values were ignored throughout. Following convergence, 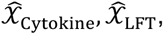 and 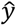 were constructed from the resulting factors and subtracted from *X*_Cytokine_, *X*_LFT_, and *y* to remove their explained covariance. Solving then proceeded to the next component, *r* + 1, using the same iterative steps above on the deflated 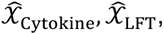 and 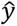 Following convergence for every factor, *D* was unstacked to recover component associations to PB and PV cytokine timepoints separately.

## Acknowledgments

This study was supported in part by National Institutes of Health grants P01AI120944 (to J.W.K.-W.) and U19AI172713 (to A.S.M.).

## Author Contributions

J.L.C., Z.C.T., J.W.K.-W., E.F.R., and A.S.M. conceived of the study. J.L.C., Z.C.T., and A.S.M. designed and performed analysis. F.M.K. collected clinical data. R.A.S., Y. Zheng, and D.W.G. managed and contributed clinical data. J.L.C., Z.C.T, and A.S.M. wrote the manuscript. R.A.S., D.W.G., A.H., F.M.K., Y. Zhai, J.W.K.-W., and E.F.R. assisted clinical interpretation and provided feedback on the manuscript.

## Data Availability

All code used to perform these analyses is available on GitHub at https://github.com/meyer-lab/liver-iri. All data used are available in this repository under the data directory at https://github.com/meyer-lab/liver-iri/tree/main/liver_iri/data.

## References

1. Lucey Michael R., Furuya Katryn N., Foley David P. Liver Transplantation. New England Journal of Medicine. 2023;389(20):1888–1900. doi:10.1056/NEJMra2200923

2. Dar WA, Sullivan E, Bynon JS, Eltzschig H, Ju C. Ischaemia reperfusion injury in liver transplantation: Cellular and molecular mechanisms. Liver Int. 2019;39(5):788–801. doi:10.1111/liv.14091

3. Wong RJ, Singal AK. Trends in Liver Disease Etiology Among Adults Awaiting Liver Transplantation in the United States, 2014-2019. JAMA Network Open. 2020;3(2):e1920294-e1920294. doi:10.1001/jamanetworkopen.2019.20294

4. Kwong AJ, Kim WR, Lake JR, et al. OPTN/SRTR 2022 Annual Data Report: Liver. American Journal of Transplantation. 2024;24(2, Supplement 1):S176–S265. doi:10.1016/j.ajt.2024.01.014

5. Deschênes M, Belle SH, Krom RAF, Zetterman RK, Lake JR, Database9 TNI of D and D and KDLT. EARLY ALLOGRAFT DYSFUNCTION AFTER LIVER TRANSPLANTATION: A Definition and Predictors of Outcome: 1. Transplantation. 1998;66(3):302.

6. Olthoff KM, Kulik L, Samstein B, et al. Validation of a current definition of early allograft dysfunction in liver transplant recipients and analysis of risk factors. Liver Transplantation. 2010;16(8):943–949. doi:10.1002/lt.22091

7. Lee DD, Singh A, Burns JM, Perry DK, Nguyen JH, Taner CB. Early allograft dysfunction in liver transplantation with donation after cardiac death donors results in inferior survival. Liver Transplantation. 2014;20(12):1447–1453. doi:10.1002/lt.23985

8. Ito T, Naini BV, Markovic D, et al. Ischemia-reperfusion injury and its relationship with early allograft dysfunction in liver transplant patients. American Journal of Transplantation. 2021;21(2):614–625. doi:10.1111/ajt.16219

9. Zhou J, Chen J, Wei Q, Saeb-Parsy K, Xu X. The Role of Ischemia/Reperfusion Injury in Early Hepatic Allograft Dysfunction. Liver Transpl. 2020;26(8):1034–1048. doi:10.1002/lt.25779

10. Selzner M, RüDiger HA, Sindram D, Madden J, Clavien PA. Mechanisms of ischemic injury are different in the steatotic and normal rat liver. Hepatology. 2000;32(6):1280–1288.

11. Upadhya AG, Topp SA, Hotchkiss RS, Anagli J, Strasberg SM. Effect of cold preservation on intracellular calcium concentration and calpain activity in rat sinusoidal endothelial cells. Hepatology. 2003;37(2). https://journals.lww.com/hep/fulltext/2003/02000/effect_of_cold_preservation_on_intracellular.13.aspx

12. Schön MR, Kollmar O, Wolf S, et al. Liver Transplantation After Organ Preservation With Normothermic Extracorporeal Perfusion. Annals of Surgery. 2001;233(1). https://journals.lww.com/annalsofsurgery/fulltext/2001/01000/liver_transplantation_after_organ_preservation.17.aspx

13. Wanner GA, Ertel W, Müller P, et al. Liver ischemia and reperfusion induces a systemic inflammatory response through Kupffer cell activation. Shock. 1996;5(1):34–40. doi:10.1097/00024382-199601000-00008

14. Hirao H, Nakamura K, Kupiec-Weglinski JW. Liver ischaemia–reperfusion injury: a new understanding of the role of innate immunity. Nature Reviews Gastroenterology & Hepatology. 2022;19(4):239–256. doi:10.1038/s41575-021-00549-8

15. Clavien PA, Camargo CAJ, Gorczynski R, et al. Acute reactant cytokines and neutrophil adhesion after warm ischemia in cirrhotic and noncirrhotic human livers. Hepatology. 1996;23(6):1456–1463. doi:10.1002/hep.510230623

16. Harmon C, Sanchez-Fueyo A, O’Farrelly C, Houlihan DD. Natural Killer Cells and Liver Transplantation: Orchestrators of Rejection or Tolerance? American Journal of Transplantation. 2016;16(3):751–757. doi:10.1111/ajt.13565

17. Obara H, Nagasaki K, Hsieh CL, et al. IFN-γ, Produced by NK Cells that Infiltrate Liver Allografts Early After Transplantation, Links the Innate and Adaptive Immune Responses. American Journal of Transplantation. 2005;5(9):2094–2103. doi:10.1111/j.1600-6143.2005.00995.x

18. Schofield ZV, Woodruff TM, Halai R, Wu MCL, Cooper MA. Neutrophils--a key component of ischemia-reperfusion injury. Shock. 2013;40(6):463–470. doi:10.1097/SHK.0000000000000044

19. Kageyama S, Nakamura K, Fujii T, et al. Recombinant relaxin protects liver transplants from ischemia damage by hepatocyte glucocorticoid receptor: From bench-to-bedside. Hepatology. 2018;68(1):258–273. doi:10.1002/hep.29787

20. Hammerich L, Heymann F, Tacke F. Role of IL-17 and Th17 Cells in Liver Diseases. Journal of Immunology Research. 2011;2011(1):345803. doi:10.1155/2011/345803

21. Liu Z, Fan H, Jiang S. CD4+ T-cell subsets in transplantation. Immunological reviews. 2013;252(1):183–191.

22. Nishimura T, Ohta A. A critical role for antigen-specific Th1 cells in acute liver injury in mice. The Journal of Immunology. 1999;162(11):6503–6509.

23. Kolda TG, Bader BW. Tensor decompositions and applications. SIAM review. 2009;51(3):455–500.

24. Acar E, Papalexakis EE, Gürdeniz G, et al. Structure-revealing data fusion. BMC Bioinformatics. 2014;15(1):239. doi:10.1186/1471-2105-15-239

25. Acar E, Kolda TG, Dunlavy DM. All-at-once optimization for coupled matrix and tensor factorizations. arXiv preprint arXiv*:11053422*. Published online 2011.

26. Chin JL, Chan LC, Yeaman MR, Meyer AS. Tensor-based insights into systems immunity and infectious disease. Trends Immunol. Published online March 29, 2023:S1471–4906(23)00047-9. doi:10.1016/j.it.2023.03.003

27. Tan ZC, Meyer AS. The structure is the message: Preserving experimental context through tensor decomposition. Cell Systems. 2024;15(8):679–693. doi:10.1016/j.cels.2024.07.004

28. Feng S, Goodrich NP, Bragg-Gresham JL, et al. Characteristics associated with liver graft failure: the concept of a donor risk index. American journal of transplantation. 2006;6(4):783–790.

29. Bro R. Multiway calibration. multilinear pls. Journal of chemometrics. 1996;10(1):47–61.

30. Van Coillie E, Van Damme J, Opdenakker G. The MCP/eotaxin subfamily of CC chemokines. Cytokine & growth factor reviews. 1999;10(1):61–86.

31. Rodríguez-Perálvarez M, Germani G, Tsochatzis E, et al. Predicting severity and clinical course of acute rejection after liver transplantation using blood eosinophil count. Transplant International. 2012;25(5):555–563.

32. Foster PF, Sankary HN, Hart M, Ashmann M, Williams JW. Blood and graft eosinophilia as predictors of rejection in human liver transplantation. Transplantation. 1989;47(1):72–74. doi:10.1097/00007890-198901000-00016

33. Lynch CA, Guo Y, Mei Z, et al. Solving the conundrum of eosinophils in alloimmunity. Transplantation. 2022;106(8):1538–1547.

34. Finkelman FD, Urban JF. The other side of the coin: The protective role of the TH2 cytokines. Journal of Allergy and Clinical Immunology. 2001;107(5):772–780. doi:10.1067/mai.2001.114989

35. Montano-Loza AJ, Rodríguez-Perálvarez ML, Pageaux GP, Sanchez-Fueyo A, Feng S. Liver transplantation immunology: Immunosuppression, rejection, and immunomodulation. Journal of Hepatology. 2023;78(6):1199–1215. doi:10.1016/j.jhep.2023.01.030

36. Duncan MD, Wilkes DS. Transplant-related Immunosuppression. Proc Am Thorac Soc. 2005;2(5):449–455. doi:10.1513/pats.200507-073JS

37. Panackel C, Mathew JF, Fawas N M, Jacob M. Immunosuppressive Drugs in Liver Transplant: An Insight. J Clin Exp Hepatol. 2022;12(6):1557–1571. doi:10.1016/j.jceh.2022.06.007

38. Ganschow R, Broering DC, Nolkemper D, et al. Th2 CYTOKINE PROFILE IN INFANTS PREDISPOSES TO IMPROVED GRAFT ACCEPTANCE AFTER LIVER TRANSPLANTATION. Transplantation. 2001;72(5). https://journals.lww.com/transplantjournal/fulltext/2001/09150/th2_cytokine_profile_in_infants_predispo ses_to.31.aspx

39. Shirwan H, Barwari L, Khan NS. PREDOMINANT EXPRESSION OF T HELPER 2 CYTOKINES AND ALTERED EXPRESSION OF T HELPER 1 CYTOKINES IN LONG-TERM ALLOGRAFT SURVIVAL INDUCED BY INTRATHYMIC IMMUNE MODULATION WITH DONOR CLASS I MAJOR HISTOCOMPATIBILITY COMPLEX PEPTIDES1,2. Transplantation. 1998;66(12). https://journals.lww.com/transplantjournal/fulltext/1998/12270/predominant_expression_of_t_helper_2_ cytokines_and.39.aspx

40. Natarajan A, Wagner B, Sibilia M. The EGF receptor is required for efficient liver regeneration. Proceedings of the National Academy of Sciences. 2007;104(43):17081–17086. doi:10.1073/pnas.0704126104

41. Komposch K, Sibilia M. EGFR Signaling in Liver Diseases. Int J Mol Sci. 2015;17(1). doi:10.3390/ijms17010030

42. Chiang EY, Kolumam G, McCutcheon KM, et al. In Vivo Depletion of Lymphotoxin-Alpha Expressing Lymphocytes Inhibits Xenogeneic Graft-versus-Host-Disease. PLOS ONE. 2012;7(3):e33106. doi:10.1371/journal.pone.0033106

43. Anders RA, Subudhi SK, Wang J, Pfeffer K, Fu YX. Contribution of the Lymphotoxin β Receptor to Liver Regeneration1. The Journal of Immunology. 2005;175(2):1295–1300. doi:10.4049/jimmunol.175.2.1295

44. Tumanov AV, Koroleva EP, Christiansen PA, et al. T Cell-Derived Lymphotoxin Regulates Liver Regeneration. Gastroenterology. 2009;136(2):694–704.e4. doi:10.1053/j.gastro.2008.09.015

45. Arend WP. The balance between IL-1 and IL-1Ra in disease. Cytokine & Growth Factor Reviews. 2002;13(4):323–340. doi:10.1016/S1359-6101(02)00020-5

46. Glimcher LH, Murphy KM. Lineage commitment in the immune system: the T helper lymphocyte grows up. Genes & development. 2000;14(14):1693–1711.

47. Kim KH, Oh EJ, Jung ES, et al. Evaluation of Pre- and Posttransplantation Serum Interferon-Gamma and Soluble CD30 for Predicting Liver Allograft Rejection. Transplantation Proceedings. 2006;38(5):1429–1431. doi:10.1016/j.transproceed.2006.03.032

48. Warlé MC, Farhan A, Metselaar HJ, et al. In vitro cytokine production of TNFα and IL-13 correlates with Acute liver transplant rejection. Human Immunology. 2001;62(11):1258–1265. doi:10.1016/S0198-8859(01)00321-4

49. Millán O, Rafael-Valdivia L, Torrademé E, et al. Intracellular IFN-γ and IL-2 expression monitoring as surrogate markers of the risk of acute rejection and personal drug response in de novo liver transplant recipients. Cytokine. 2013;61(2):556–564. doi:10.1016/j.cyto.2012.10.026

50. Bockhorn M, Goralski M, Prokofiev D, et al. VEGF is important for early liver regeneration after partial hepatectomy. J Surg Res. 2007;138(2):291–299. doi:10.1016/j.jss.2006.07.027

51. Yamamoto C, Yagi S, Hori T, et al. Significance of Portal Venous VEGF During Liver Regeneration After Hepatectomy. Journal of Surgical Research. 2010;159(2):e37–e43. doi:10.1016/j.jss.2008.11.007

52. Yao J, Feng XW, Yu XB, et al. Recipient IL-6-572C/G genotype is associated with reduced incidence of acute rejection following liver transplantation. J Int Med Res. 2013;41(2):356–364. doi:10.1177/0300060513477264

53. Karimi MH, Daneshmandi S, Pourfathollah AA, et al. Association of IL-6 promoter and IFN-γ gene polymorphisms with acute rejection of liver transplantation. Molecular Biology Reports. 2011;38(7):4437–4443. doi:10.1007/s11033-010-0572-6

54. García de la Garza R, Sarobe P, Merino J, et al. Immune monitoring of immunosuppression withdrawal of liver transplant recipients. Transplant Immunology. 2015;33(2):110–116. doi:10.1016/j.trim.2015.07.006

55. Sosa RA, Zarrinpar A, Rossetti M, et al. Early cytokine signatures of ischemia/reperfusion injury in human orthotopic liver transplantation. JCI Insight. 2016;1(20):e89679. doi:10.1172/jci.insight.89679

56. Pedregosa F, Varoquaux G, Gramfort A, et al. Scikit-learn: Machine learning in Python. the Journal of machine Learning research. 2011;12:2825–2830.

57. Davidson-Pilon C. lifelines: survival analysis in Python. Journal of Open Source Software. 2019;4(40):1317.

58. Roald M, Moe YM. TLViz: Visualising and analysing tensor decomposition models with Python. Published online 2022.

59. Virtanen P, Gommers R, Oliphant TE, et al. SciPy 1.0: fundamental algorithms for scientific computing in Python. Nat Methods. 2020;17(3):261–272. doi:10.1038/s41592-019-0686-2

